# Explainable and Calibrated AI for Decoding Host-Adaptive Changes in Influenza A Virus

**DOI:** 10.64898/2026.05.23.726879

**Authors:** Hoang-Hai Nguyen, Josip Rudar, Samira Mubareka, David Lapen, Yohannes Berhane, Carson K. Leung, Oliver Lung

**Affiliations:** National Centre for Foreign Animal Disease, Canadian Food Inspection Agency, Winnipeg, Manitoba R3E 3M4, Canada; Department of Computer Science, University of Manitoba, Winnipeg, Manitoba R3T 2N2, Canada; Sunnybrook Research Institute, Toronto, Ontario M4N 3M5, Canada; Department of Laboratory Medicine & Pathobiology, University of Toronto, Toronto, Ontario M5S 1A8, Canada; Science and Technology Branch, Agriculture and Agri-Food Canada, Ottawa, Ontario K1A OC6, Canada; Department of Biological Sciences, University of Manitoba, Winnipeg, Manitoba R3T 2N2, Canada

**Keywords:** deep learning, explainable AI, host adaptation, influenza A virus

## Abstract

**Background:** Influenza A virus (IAV) is a major public health burden, causing seasonal epidemics and occasional pandemics. Its transmission from avian species to mammals and subsequent spread requires adaptive changes in the viral genome. Understanding these molecular adaptations is essential for pandemic preparedness, and machine learning offers a powerful approach to uncover the evolution and biology of IAV.

**Results:** This study established a well-calibrated WaveSeekerNet model that accurately predicted the host source across all 8 IAV segments (macro F1-score: 0.9728), significantly improving the reliability of predicted probabilities with calibration errors approaching zero. Model interpretation revealed that avian-adapted IAVs consistently activated G/C content, whereas mammalian-adapted IAVs generally activated A/T content. This distinction was confirmed by codon-level analysis, in which G/C-rich codons were rewarded for the avian hosts and A/T-rich codons for the mammalian hosts. In the feature space learned by WaveSeekerNet, we defined host-adaptive distance to quantify species barriers and proposed it as a risk-assessment metric. We hypothesized the Mammalian Adaptation Zone (MAZ), a zone where the virus is expected to adjust its host-adaptive distance to reach, thereby helping it establish persistent mammalian lineages. The analysis also revealed the Hard Distance of avian-origin viruses (e.g., H5Nx, H9N2), indicating they have not yet established persistent mammalian lineages. Finally, analysis of human H7N9 (2013, China) and non-human mammalian H5Nx (North America) viruses showed that WaveSeekerNet accurately identified key mammalian-adaptive mutations, including PB2-E627K and PB2-D701N.

**Conclusions:** WaveSeekerNet elucidated IAV host-adaptation mechanisms in silico, providing insights into the underlying mechanisms of host adaptation and informing improved surveillance and intervention strategies.

## 1 Introduction

Influenza A virus (IAV) is an enveloped, negative-sense, single-stranded RNA virus with a genome of 8 segments—Segment 1 (PB2), Segment 2 (PB1), Segment 3 (PA), Segment 4 (HA), Segment 5 (NP), Segment 6 (NA), Segment 7 (M) and Segment 8 (NS)—and is classified within the *Orthomyxoviridae* family [1]. IAV naturally circulates among waterfowl, frequently causing spillover infections in mammalian hosts, including swine, horses, dogs, and humans [2,3]. Understanding the mechanisms underlying IAV’s broad host range is critical, as this virus has caused multiple pandemics/panzootics and ongoing outbreaks in humans, livestock, and wildlife, leading to substantial economic losses and mortality. For example, since 2021, there has been an ongoing outbreak of highly pathogenic avian influenza (HPAI) caused by IAV from clade 2.3.4.4b. This HPAI virus has led to unprecedented outbreaks in wild birds, domestic poultry, and dairy cows [4,5], resulting in severe livestock mortality events and major economic losses for farmers. Historically, IAV has exhibited a remarkable capacity for zoonotic spillover, driving 4 major pandemics since 1918—the Spanish flu 1918 (H1N1), the Asian flu 1957 (H2N2), the Hong Kong flu 1968 (H3N2), and the 2009 swine flu pandemic (H1N1) [6]—as well as recurrent seasonal epidemics that remain a persistent global public health burden. Therefore, understanding the molecular determinants of host adaptation and efficient mammal-to-mammal transmission contributing to IAV’s ability to breach species barriers is urgently needed to better prepare for future outbreaks and pandemics.

In recent years, the integration of Machine Learning (ML) and Artificial Intelligence (AI)—particularly through explainable AI methods—has opened new avenues for exploring complex biological phenomena such as IAV host adaptation [7–10]. These computational approaches can uncover subtle yet critical patterns in IAV’s vast genomic sequence data that traditional sequence-based analyses may overlook. Although ML and AI methods can identify highly impactful mutations or positions with mechanistic implications, most existing studies focus on predictive tasks—such as host preferences or transmission risks [8,10–13]—rather than dissecting the mechanistic basis for these predictions and how they contribute to IAV host adaptation. However, modern deep learning models often exhibit miscalibration [14,15], leading to unreliable predictions that hinder their application in high-stakes domains.

In this study, we establish a novel framework based on our recently developed WaveSeekerNet model—a classifier trained to accurately predict IAV subtypes and host source [16]—to deliver reliable host-source predictions for IAV while providing mechanistic insights into host adaptation. We used SHapley Additive exPlanations (SHAP) [17,18] to systematically analyze large-scale IAV genomic sequences and identify patterns that likely govern host adaptation and efficient mammal-to-mammal transmission. Nucleotide- and codon-level SHAP analyses revealed that avian-adapted viruses exhibit elevated G/C content, whereas mammalian-adapted strains show A/T enrichment. We further defined a host-adaptive distance metric to characterize IAV transitions within host-adaptive space, proposing the concepts of the “Mammalian Adaptation Zone” (MAZ) and the “Hard Distance.” These concepts suggest that avian-origin viruses (e.g., H5Nx, H9N2) have not yet established persistent mammalian lineages.

We also proposed host-adaptive distance as a risk-assessment metric for monitoring IAV evolution. In addition, we demonstrated WaveSeekerNet’s ability to identify key adaptive mutations (PB2-E627K and PB2-D701N) associated with host adaptation. These findings provide new insights into IAV evolution and biology, informing resource allocation and strategy in future surveillance and intervention programs.

## 2 Materials and Methods

### 2.1 Influenza A virus dataset

We downloaded the RNA sequences of IAV whole genomes from GISAID EpiFlu in September 2025, along with subtype, host, collection date, and other available metadata [19]. The data preparation workflow is illustrated in Fig. 1A. To minimize bias in model training and downstream analysis, we removed duplicate sequences and retained only the earliest collected sequences. We filtered sequences by length and ambiguity, retaining only those whose length differed by no more than 20% from that of the reference strain *A/New York/392/2004* (H3N2) and that contained at most 10 ambiguous bases. We further screened and annotated sequences using the VADR (Viral Annotation DefineR) tool [20]. Sequence validation and annotation were performed using the “v-annotate.pl” VADR script (v1.6.4) with the influenza models (v1.6.3–2) [21] and default parameters, retaining only sequences that passed VADR filtering and contained protein coding sequence annotations in the final dataset. We grouped IAV sequences into three major host categories: humans, avian (e.g., falcons, turkeys, goose), and non-human mammals (e.g., swine, horses, dogs). Finally, to mimic real-world forecasting scenarios where models are trained on historical data to predict future events, we implemented a temporal data split. Sequences collected before January 1, 2020, were used for training, while those collected from 2020 onwards were reserved for model evaluation. The final dataset comprises 1,292,288 sequences of 8 segments, including 690,773 pre-2020 and 601,515 post-2020 sequences. To support our analysis, we also prepared a separate dataset of IAV H5Nx sequences from ongoing North American outbreaks, comprising 68,295 sequences for all 8 segments. Data distribution details are provided in Additional file 2: Supplementary Table S1.

**Figure 1.**
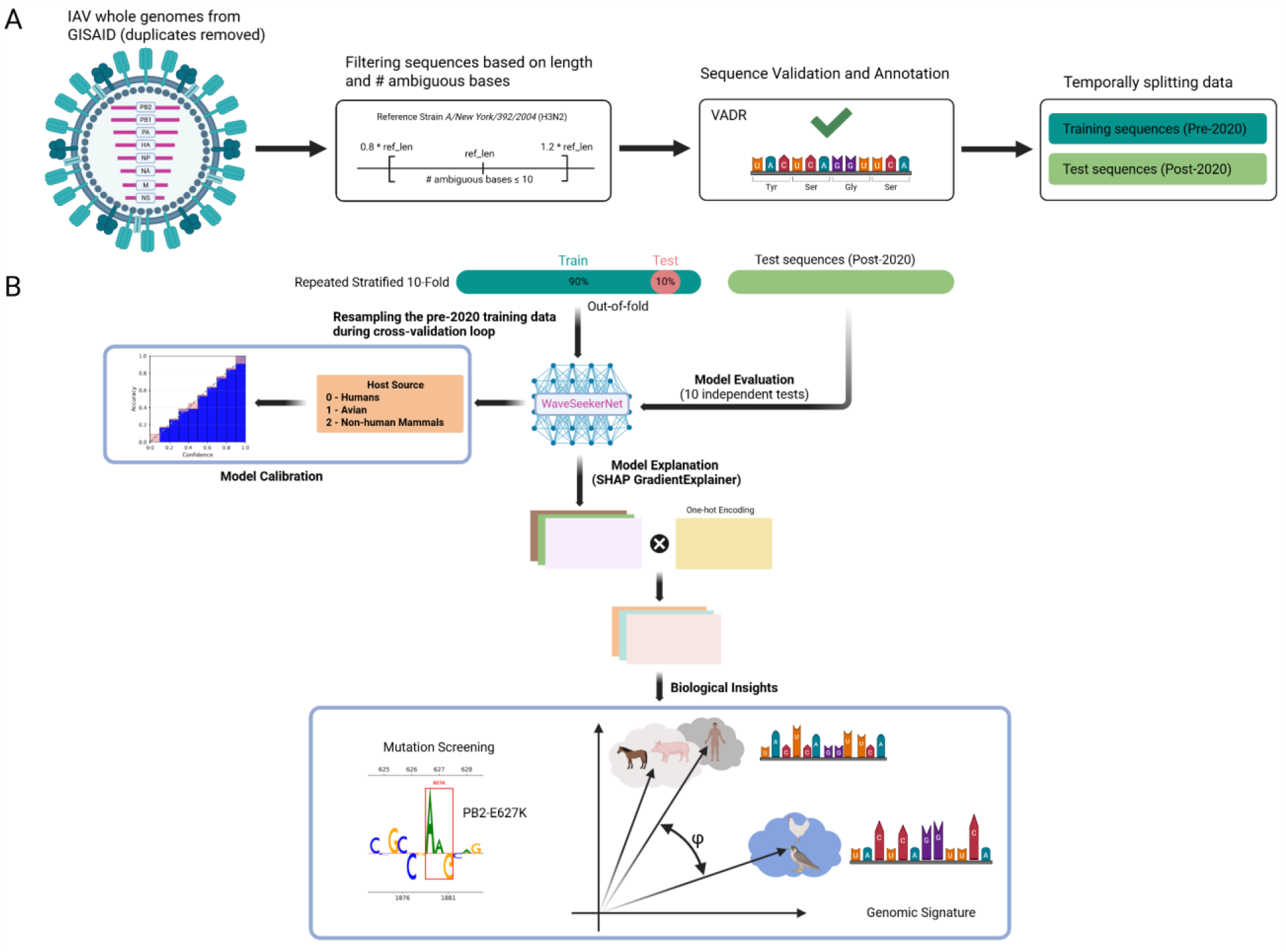
Illustrations of data processing, model training, and downstream analysis. (A) Whole-genome IAV sequences were retrieved from the GISAID EpiFlu database along with associated metadata, including subtype, host, and collection date. Duplicate sequences were removed based on the collection date, retaining the earliest collected record. The remaining sequences underwent quality control and validation/annotation using VADR. Sequences that passed VADR validation were temporally split into pre-2020 training and post-2020 test datasets. (B) The WaveSeekerNet model was trained using repeated stratified 10-fold cross-validation, with resampling performed within the cross-validation loop. After training, to obtain reliability of predicted probabilities, we performed model calibration using isotonic regression. SHAP values were computed for each sequence using GradientExplainer. These values were then multiplied element-wise by the corresponding one-hot-encoded sequences to derive the actual nucleotide-level attribution scores, which were used to inform biological insights. The figure was created with BioRender, in which the reliability diagram was generated using Python code with mock data representing idea calibration.

### 2.2 The WaveSeekerNet model

We recently introduced a deep-learning model, WaveSeekerNet [16], which was used to analyze IAV and predict subtypes and the host source. Briefly, this model is a transformer-based architecture that employs an ensemble of attention-like mechanisms to identify patterns in IAV genomes and use these patterns for classification. The key component of WaveSeekerNet is the WaveSeeker block, which comprises an ensemble of 3 attention-like mechanisms: the Fourier Transform, the Wavelet Transform, and the gated multi-layer perceptron (gMLP) [22]. WaveSeekerNet first splits the image representation of sequences, such as the Frequency Chaos Game Representation (FCGR) or One-hot encoding, into distinct “word” patches, known as tokens. Transformed tokens are then analyzed by the WaveSeeker block before being passed to the Global Expectation Pooling for dimension reduction [23], and finally to the Classification Head for classification.

### 2.3 Model training and evaluation

To enable training of the WaveSeekerNet model, we converted RNA sequences to one-hot encoding, with each character represented as a 5-dimensional vector. In this study, adenine (A) was encoded as (1, 0, 0, 0, 0), cytosine (C) as (0, 1, 0, 0, 0), guanine (G) as (0, 0, 1, 0, 0), and thymine (T) as (0, 0, 0, 1, 0). All ambiguous bases (e.g., N, Y, R, W) were mapped to N and encoded as (0, 0, 0, 0, 1). Zero-padding was also performed to ensure all model inputs had the same length. Table 1 presents the available settings for the WaveSeekerNet model; in this study, we used the baseline configuration with all model settings enabled. The model was trained on a high-performance computing cluster equipped with NVIDIA A100 GPUs.

**Table 1:**
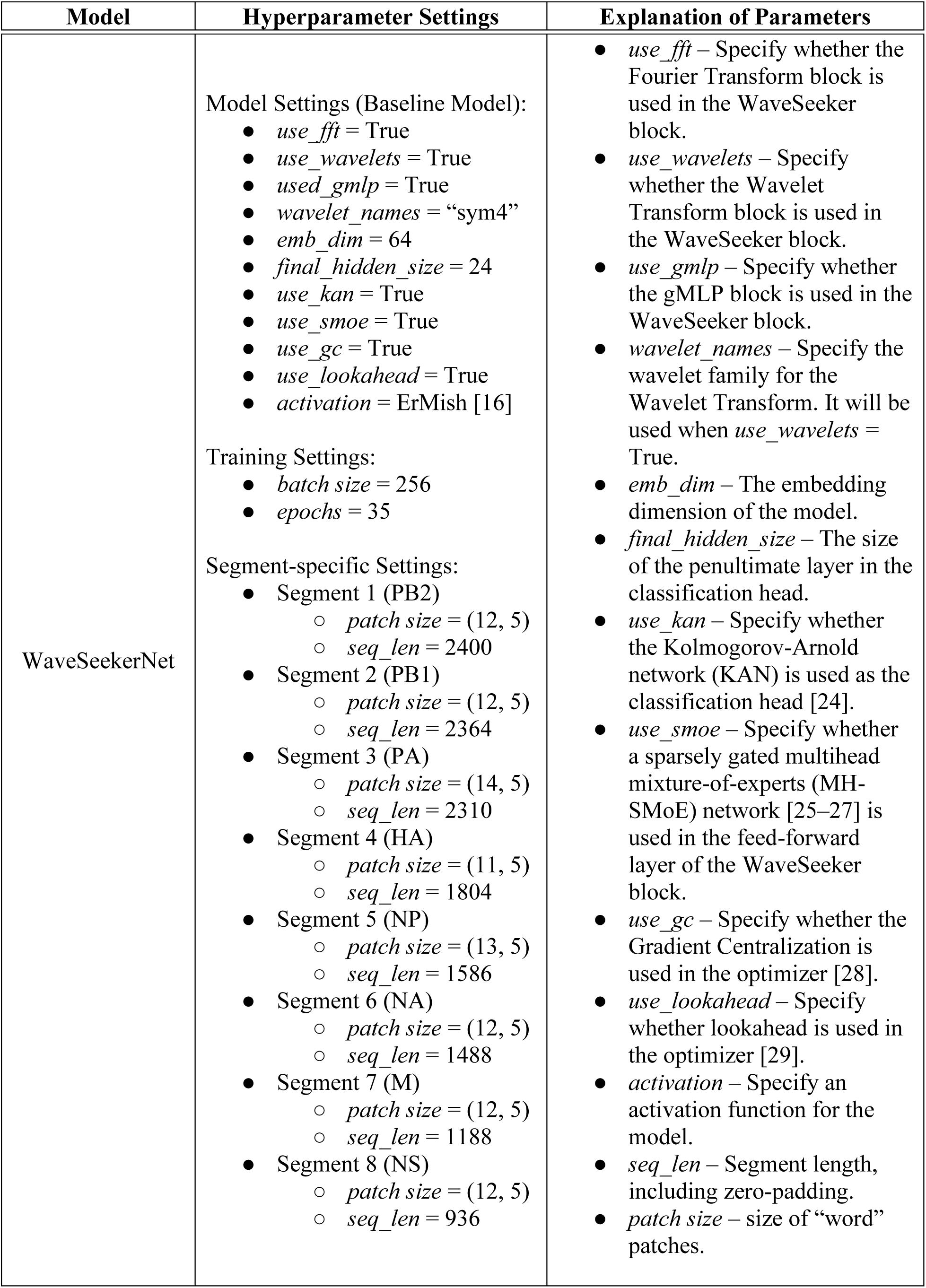
The hyperparameter settings used to train the WaveSeekerNet model.

The dataset shows class imbalance, with human and avian sequences comprising the majority of the data. To prevent bias during model training, we downsampled human and avian sequences to 16,000 each while retaining all non-human mammalian sequences. We performed random downsampling using the “resample” function from scikit-learn (v.1.5.1). We applied the same training strategy as in our previous study [16], using repeated stratified 10-fold cross-validation (*n_repeats* = 1) with resampling of pre-2020 training data at each iteration of the cross-validation loop (Fig. 1B). Evaluation metrics include F1-score (Macro Average), Balanced Accuracy (BA), and Matthews Correlation Coefficient (MCC). Final model evaluation includes out-of-fold pre-2020 data and 10-fold post-2020 data, covering the entire dataset (Fig. 1B).

### 2.4 Model calibration

Deep neural networks (DNNs) often display overconfidence and/or underconfidence in their predictions [14,15]. This miscalibration means predicted probabilities do not accurately reflect the true likelihood of correctness (e.g., showing overconfidence in errors or underconfidence when correct). Calibration methods, applied post-training or integrated into the training objective, adjust output probabilities to yield reliable estimates. These are crucial for trustworthy uncertainty quantification in high-stakes applications. In this study, we applied isotonic regression to calibrate WaveSeekerNet’s probability outputs (Fig. 1B). To handle multiple classes, we adopted a one-versus-all strategy, decomposing the multiclass problem into binary subproblems, calibrating each via isotonic regression, and aggregating the resulting probabilities [30]. To assess whether isotonic regression performs well on our dataset, we performed isotonic calibration using 5-fold cross-validation on the pre-2020 and post-2020 datasets separately. For the post-2020 dataset, we repeated the 5-fold cross-validation procedure 10 times to align with the model-training strategy’s 10-fold structure (10 independent tests; Fig. 1B). For each calibration procedure, we trained a calibrator on four folds and applied it to the held-out fold. This produced out-of-fold calibrated probabilities for all training samples, preventing overfitting and providing an unbiased estimate of calibration performance.

To comprehensively evaluate calibration performance, we employed a suite of complementary metrics—Expected Calibration Error (ECE) [31], Maximum Calibration Error (MCE) [31], Miscalibration Score (MCS) [15], and Adaptive Calibration Error (ACE) [32]—alongside reliability diagrams for visual assessment [14]. ECE partitions predictions into *M* fixed-width bins and calculates a weighted average of the absolute difference between accuracy and confidence (predicted probability) across all bins:

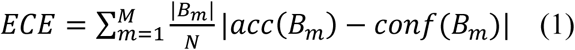

where *N* is the total number of samples, 𝐵_𝑚_ is the set of samples whose prediction confidence falls into the interval of bin *m*, |𝐵_𝑚_| is the number of samples in bin *m*, 𝑎𝑐𝑐(𝐵_𝑚_) is the average accuracy of samples in bin *m*, and 𝑐𝑜𝑛𝑓(𝐵_𝑚_) is the average confidence (predicted probability) of samples in bin *m*. In multi-class scenarios, the standard ECE typically considers the top-1 prediction (the class with the highest probability). MCE captures the worst bin, where the gap between confidence and accuracy is largest:

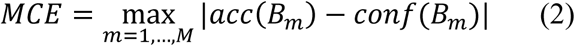

Unlike ECE’s absolute gaps, MCS preserves sign to diagnose miscalibrated directionality: MCS > 0 indicates overconfidence; MCS < 0 reveals underconfidence:

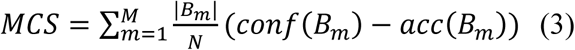

Standard ECE uses fixed-width bins (e.g., 0.0–0.1, 0.1–0.2, …, 0.9–1.0). Overconfident models concentrate samples in the 0.9–1.0 bin, leaving earlier bins empty and biasing ECE toward high-confidence regions. ACE addresses this via equal-frequency (mass) binning, dividing sorted confidences into *R* bins of ∼*N/R* samples each. This ensures calibration error reflects the full distribution of model outputs, not just high-density regions:

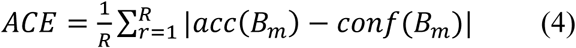

### 2.5 SHAP features attribution and codon usage bias

SHAP values provide a standardized metric for feature importance, quantifying each feature’s individual contribution to a prediction. We estimated SHAP values using GradientExplainer implemented in the SHAP package (v0.48.0) [17]. The SHAP GradientExplainer combines Shapley value theory with the computational efficiency of Expected Gradients—an extension of Integrated Gradients [33]—to approximate feature attributions for each nucleotide position in the sequence. This computationally efficient approximation is well-suited for computing SHAP values across 3 host classes for more than 1.3 million sequences in our dataset, using a high-performance computing cluster equipped with NVIDIA A100 GPUs.

We computed SHAP values for all 3 classes (0 - Humans, 1 - Avian, and 2 - Non-human Mammals) for each sequence, capturing feature attributions for the ground-truth label as well as the other 2 classes (see Fig. 1B). To reduce SHAP GradientExplainer computation time while maintaining sufficient accuracy, we selected 1,000 training sequences as background data—350 sequences each for human and avian hosts and 300 for non-human mammalian hosts—and used a batch size of 256. With this setting, the computation can take up to 4 days for 10 folds of the HA segment alone (105,804 sequences) in the post-2020 dataset. Final SHAP values for downstream analysis included out-of-fold pre-2020 data, 10-fold post-2020 data, and H5Nx North American data, totaling 1.5 terabytes (TB) of SHAP values stored in NumPy format. For each training fold, we computed SHAP values for the 61 sense codons in every sequence of the post-2020 dataset. The 10 resulting sets of codon-level SHAP values (10 independent tests; Fig. 1B) were averaged to mitigate overfitting, and the resulting averages were used for downstream analysis. To visualize the SHAP values for each codon nucleotide as a sequence logo of model importance scores, we used the “viz_sequence.py” script from the DeepLIFT package [34].

In this study, we also examined whether codon-SHAP values are related to codon usage bias. Relative Synonymous Codon Usage (RSCU) is a widely used metric to quantify codon usage bias, measuring how frequently a specific codon is used relative to its expected frequency if all synonymous codons for that amino acid were used with equal probability [35]. RSCU > 1 indicates the codon is used more frequently than expected, RSCU = 1 indicates no bias, and RSCU < 1 indicates the codon is used less frequently than expected. To quantify the extent to which codon usage bias aligns with codon-level model importance, we calculated the Spearman correlation coefficient 𝜌 and corresponding p-values between RSCU values and SHAP values for each individual codon. The resulting p-values were adjusted for multiple testing using the Benjamini–Hochberg procedure to control the false discovery rate (q-values; FDR control).

### 2.6 Definition of host-adaptive distance

The WaveSeekerNet model acts as a function that maps the tokenized representation of each IAV segment to its phenotype (e.g., human hosts). After training, SHAP can be used to compute attribution scores that quantify the contribution of each nucleotide in the tokenized input to the model’s final prediction. Consequently, sequences from different host groups may occupy distinct regions in a multi-dimensional space, reflecting host-specific patterns learned by the WaveSeekerNet model (Fig. 1B). We adapt the SHAP distance methodology [36] to define host-adaptive distance as the directional dissimilarity in functional reasoning captured by the WaveSeekerNet model. A feature attribution vector of SHAP values for the 61 sense codons, Φ*_X_* = 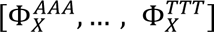, is calculated for each sequence. The host-adaptive distance between two sequences, *A* and *B*, can then simply be calculated using the cosine distance:

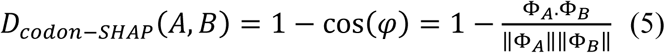

where 𝜑 is the angle between the two feature attribution vectors Φ_𝐴_ and Φ_𝐵_.

The value of *D* lies between 0 and 2. When *D* = 0, the feature attribution vectors of two sequences share the same trajectory (identical). When *D* = 1, they are orthogonal. When *D* = 2, they point in exactly opposite directions. The cosine distance is particularly appropriate for studying convergent evolution and host adaptation because it remains invariant to differences in sequence length and codon usage bias while capturing directional dissimilarity in trajectories. Intuitively, one can think of this as a comparison between two documents on the same topic, each with a different word count. The cosine distance correctly identifies that these documents share the same topic despite their different lengths.

## 3 Results

### 3.1 The calibrated WaveSeekerNet model correctly identifies the host source

In this analysis, we focused on two key aspects of model performance: the model’s ability to make correct predictions (discriminative accuracy) and the reliability of its associated confidence scores (probabilistic calibration). The analysis results for post-2020 data are summarized in Figs. 2, 3, and Additional file 1: Supplementary Fig. S1. Seven segments (PB2, PB1, PA, NP, NA, M, and NS) achieved F1-scores (Macro Average) ranging from 0.9229 ± 0.0169 to 0.9580 ± 0.0114, MCC values ranging from 0.9398 ± 0.0152 to 0.9615 ± 0.0059, and BA values ranging from 0.9579 ± 0.0010 to 0.9713 ± 0.0012 (pink bars in Fig. 2). The discriminative ability of segment HA was notably lower, achieving an F1-score (Macro Average) of 0.8839 ± 0.0672, an MCC of 0.8370 ± 0.1068, and a BA of 0.9478 ± 0.0102.

**Figure 2.**
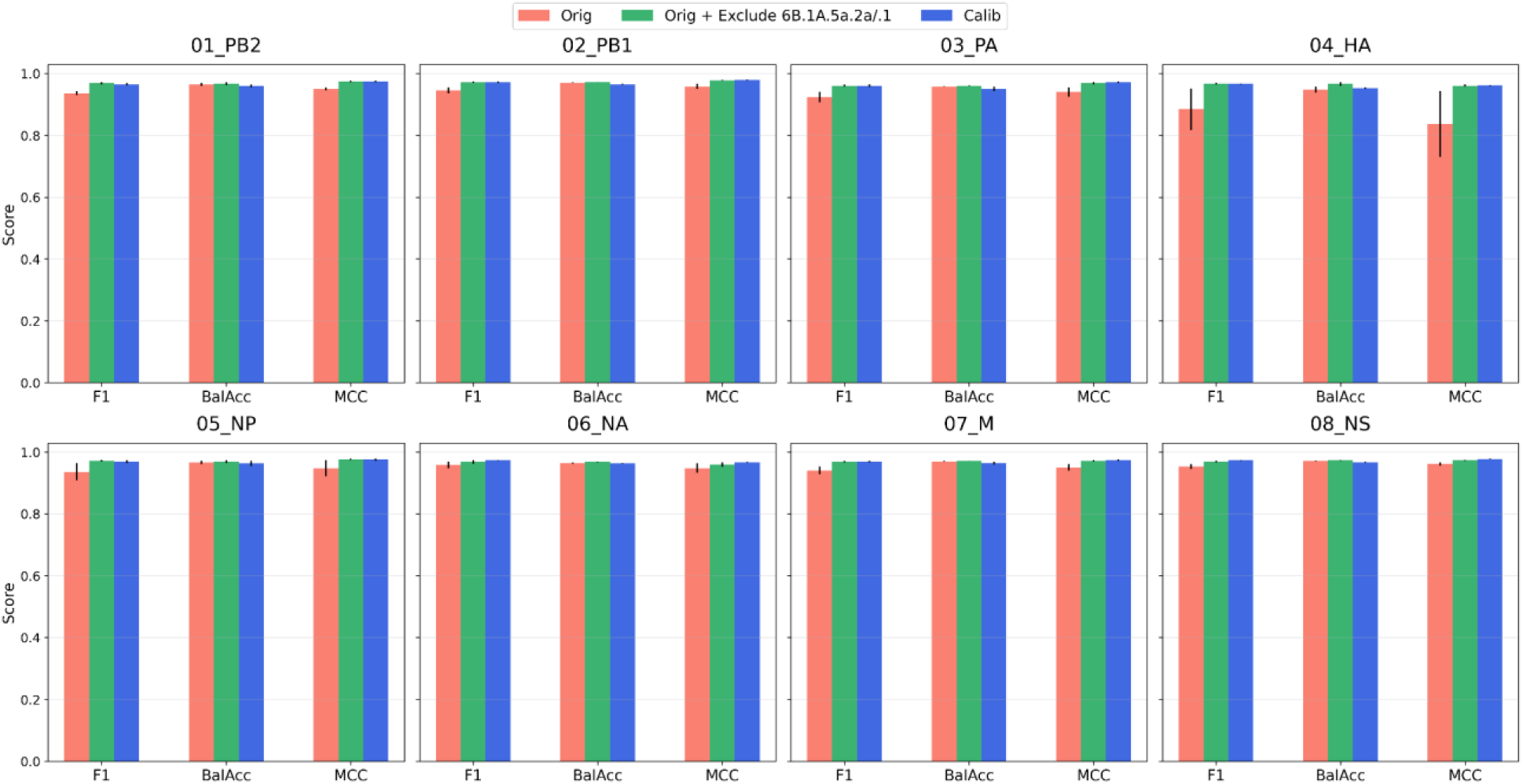
Model performance on the post-2020 dataset. Bar colors indicate: pink, before calibration; green, before calibration with human H1N1 seasonal viruses from clades 6B.1A.5a.2a and 6B.1A.5a.2a.1 excluded; blue, after calibration. Performance is evaluated using F1-score (Macro Average), BA, and MCC.

**Figure 3.**
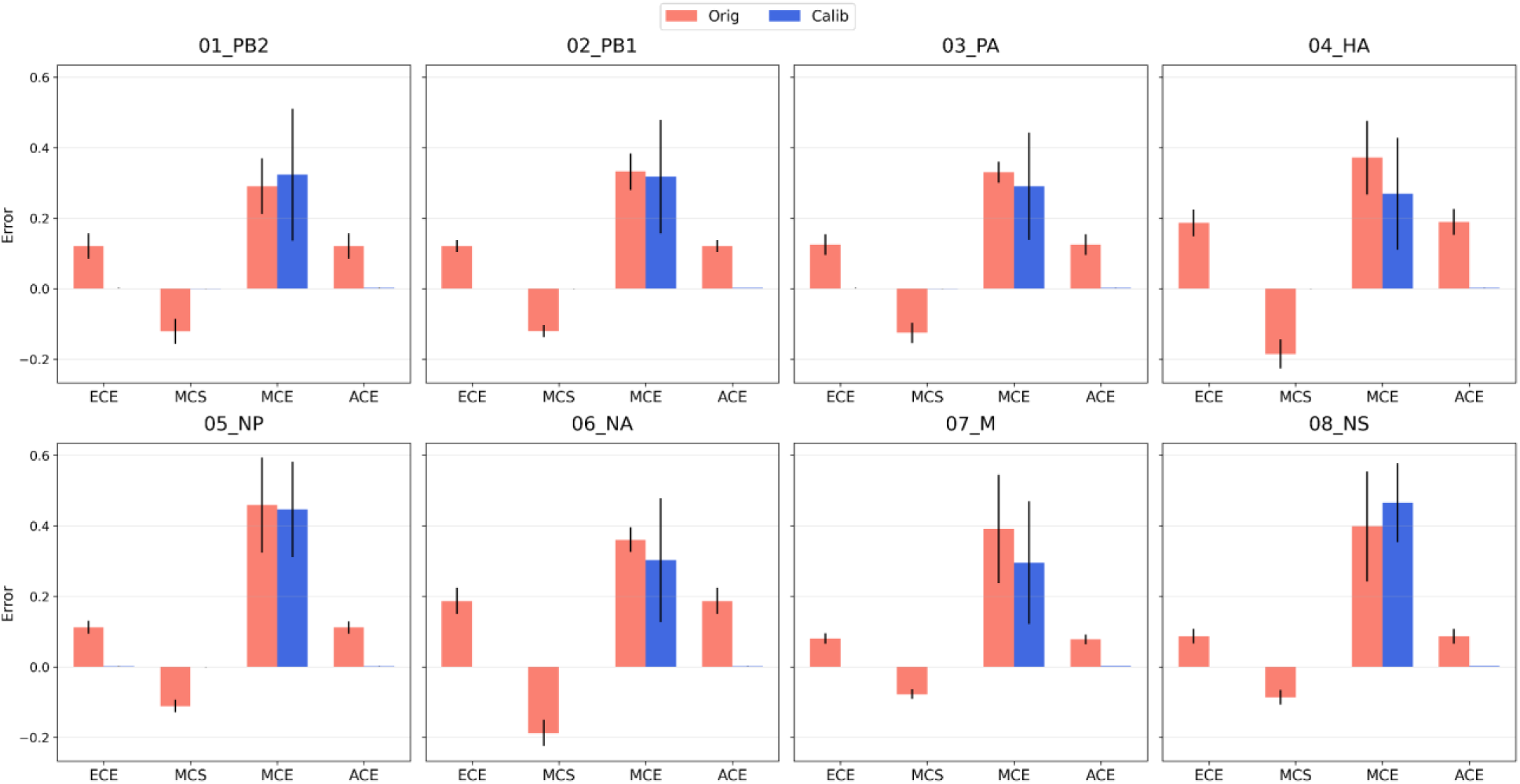
Calibration results for the post-2020 dataset. Calibration is evaluated using ECE, MCS, MCE, and ACE.

We further analyzed the reliability of the model’s confidence scores. Fig. 3 and Additional file 1: Supplementary Fig. S1 show calibration results and reliability diagrams, respectively. The uncalibrated (pink) curves (Additional file 1: Supplementary Fig. S1) for all 8 segments consistently lie above the diagonal line of perfect calibration alongside negative MCS (Fig. 3), indicating underconfidence. Before calibration, mean ECE and ACE values across all 8 segments were relatively high, roughly ranging from 0.09 to 0.2, with the largest miscalibration occurring with segment HA (ECE and ACE values were 0.1872 ± 0.0380, 0.1893 ± 0.0369, respectively), indicating a substantial deviation between predicted confidence and empirical accuracies (pink bars in Fig. 3). However, after calibration, the ECE and ACE values were improved significantly for all 8 segments, with reductions of at least 99.1% (blue bars in Fig. 3). For example, on segment HA, the ECE value decreased from 0.1872 ± 0.0380 to 0.0003 ± 0.0001, a 99.8% reduction. In terms of discriminative accuracy (blue bars in Fig. 2), across all 8 segments, the calibrated models achieved F1-scores (Macro Average) ranging from 0.9606 to 0.9728, with very low variance (±0.0006 to 0.0049). The MCC values consistently exceeded 0.9608, with the highest performance observed in NS (0.9770 ± 0.0008) and PB2 (0.9783 ± 0.0016). These values indicated that calibration either maintained or improved the model’s ability to correctly distinguish between avian, human, and non-human mammalian hosts. We also reported the MCE values, which represent the worst-case bin. Although the MCE values were relatively high, up to 0.4660, the number of samples falling into the worst bins (the bins with the large gap) was very small across all 8 segments, with a maximum of 1% of samples being poorly calibrated when their gaps exceeded 0.05 (see the reliability diagram of segment PB2 in Additional file 1: Supplementary Fig. S1). Additional file 1: Supplementary Figs. S2, S3, and S4 present the analyses of pre-2020 data, revealing patterns similar to those observed in post-2020 data. Together, analyses of both datasets demonstrated that isotonic regression dramatically improved the reliability of the probability estimates while maintaining predictive accuracy.

### 3.2 The impact of H1N1 seasonal viruses on WaveSeekerNet’s predictive performance

Observed that the model’s predictive performance was not particularly high across all segments before calibration. Thereupon, we investigated why the model did not achieve the expected high performance and found that human H1N1 seasonal viruses belonging to clades 6B.1A.5a.2a and 6B.1A.5a.2a.1—the subclades of A(H1N1)pdm09 [37]—were the cause. We observed that only a few H1N1 sequences (fewer than 6 per segment) in these clades were present in the pre-2020 training data (Additional file 2: Supplementary Table S1); therefore, we assessed their impact only on the post-2020 dataset. Since 2024, these sequences have accounted for the majority of submissions to the EpiFlu GISAID database. They comprised 40,308 (38.1%) of the 105,804 post-2020 HA sequences in our dataset. Examining the failure details across 10 training folds for segment HA, we found that many of these human sequences were misclassified as non-human mammals. For example, in the worst case (training fold 4), the model misclassified 12,274 HA sequences of human H1N1 seasonal viruses as non-human mammals—9,356 submitted in 2024–2025 and 2,918 in 2022–2023—yielding the lowest F1-score (Macro Average) of 0.7794. To evaluate the impact of sequences from clades 6B.1A.5a.2a and 6B.1A.5a.2a.1, we assessed the model’s predictive performance on the post-2020 dataset (uncalibrated) after their exclusion. The model then achieved high and stable performance across 10 folds for all 8 segments (green bars in Fig. 2), with F1-scores (Macro Average) ranging from 0.9599 ± 0.0033 to 0.9721 ± 0.0028. Segment HA achieved an F1-score (Macro Average) of 0.9674 ± 0.0029, compared with 0.8839 ± 0.0672 when these sequences were included in the post-2020 dataset. We further performed class-wise calibration analyses to explain this misclassification between human and non-human mammalian classes (Additional file 3: Supplementary Table S2). The human class exhibited underconfidence (MCS < 0), while the non-human mammalian class showed overconfidence (MCS > 0) across segments and 10 training folds. This happened when the model assigned low probabilities to the correct human class and high probabilities to the non-human mammalian class, even to sequences that are not truly non-human mammalian (False Positive). Interestingly, the avian class showed low mean ECE, ACE, and absolute MCS < 0.02 before calibration, indicating the model’s reliable confidence in distinguishing avian from mammalian sources.

### 3.3 The significance of nucleotide composition in the host adaptation of IAV

To elucidate the fundamental mechanisms of IAV host adaptation, we computed compositional SHAP values for each nucleotide type across all segments. This approach allows us to pinpoint nucleotide types that significantly influence the model’s predictions. The results for the post-2020 and pre-2020 data are shown in Fig. 4 and Additional file 1: Supplementary Fig. S5, respectively. The G/C content was a consistently strong positive predictor for avian hosts, with positive SHAP values observed across all 8 segments (Fig. 4, Additional file 1: Supplementary Fig. S5). Conversely, A/T content had neutral to negative predictive impacts (near-zero or negative SHAP values). The uniform G/C bias across segments underscores nucleotide composition as a consistent biological signature that fundamentally shapes the model’s avian host predictions.

**Figure 4.**
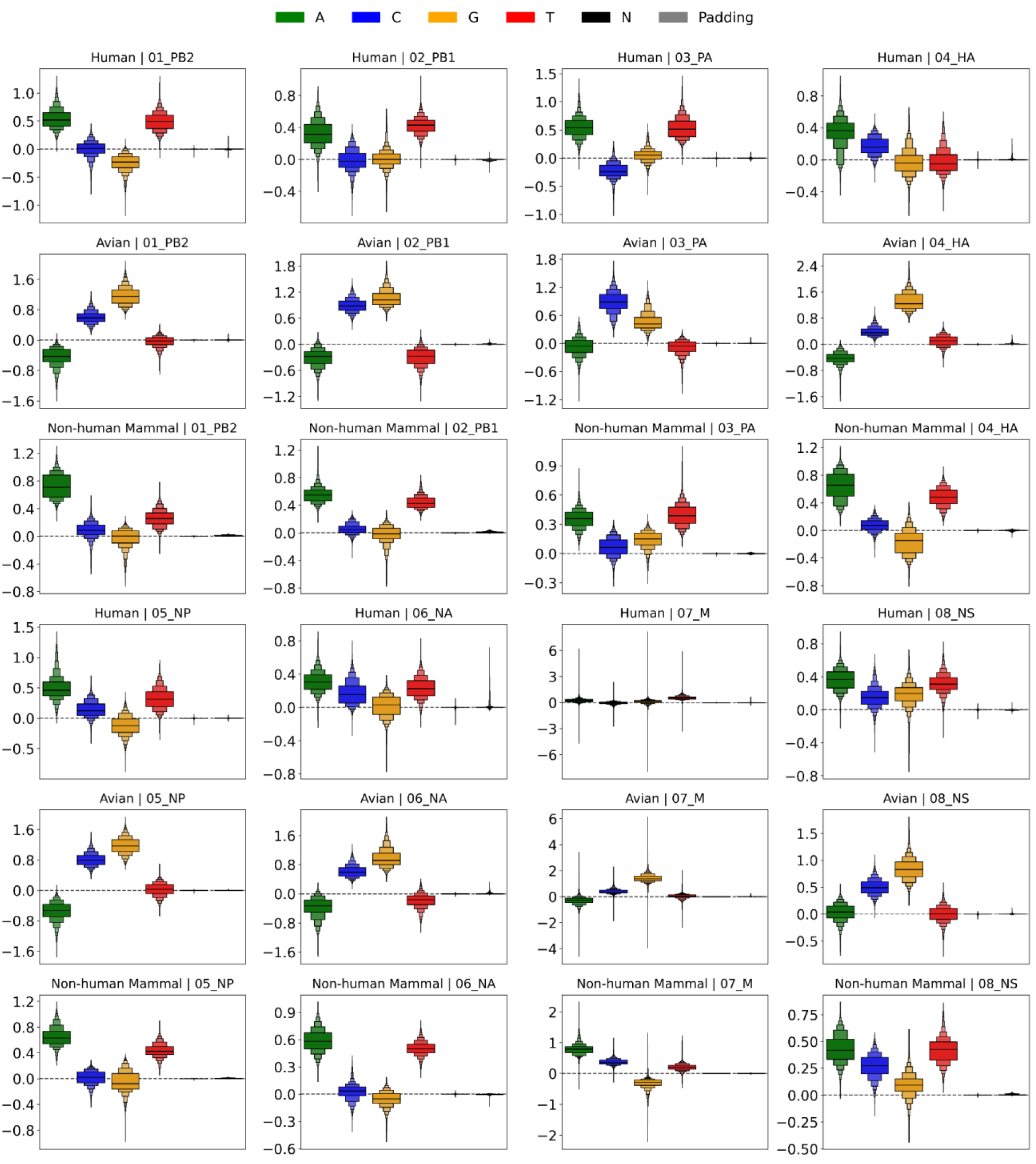
Distribution of nucleotide-SHAP values across host groups and genome segments on the post-2020 dataset. Boxen plots show SHAP value distributions for correctly predicted sequences, aggregated across 10 cross-validation folds.

In contrast, non-human mammalian hosts generally showed the opposite trend, with elevated positive SHAP values for A/T content and near-zero to negative values for G/C across six segments (PB2, PB1, PA, HA, NP, and NA). Segments M and NS also showed positive SHAP contributions for C content. Human hosts had a similar signature, with higher positive SHAP values associated with A/T content. However, we observed modest positive contributions to predictions from certain G/C content—C in segments HA, NP, and NA; G in segment M; and both G and C in segment NS—in human-derived sequences, though these contributions remained consistently lower than the observed A/T signals. We observed outliers in training fold 10, which affected the scale of the visualization for segment M but did not affect the interpretation of the results. Finally, we observed that neither zero-padding from one-hot encoding nor ambiguous bases affected the model’s predictions, as they had nearly zero SHAP values, demonstrating that the model effectively learned functional patterns within sequences and used these patterns for predictions.

The model achieved high performance (macro F1-score: 0.9728), indicating robust classification, although a few sequences were misclassified. When misclassification occurred, it generally involved sequences with a host-source discrepancy. Our previous work demonstrated that this is indicative of a biologically significant event where the virus retains its original host signature after cross-species transmission [16]. As shown in Additional file 1: Supplementary Figs. S6 and S7, the SHAP analysis revealed that the distribution of feature importance for misclassified sequences was consistent with that for correctly classified sequences. This consistency suggests that the model’s decisions, even when incorrect, were based on recognizing stable, biologically meaningful functional patterns, confirming that the model genuinely learned relevant features rather than relying on noise.

### 3.4 WaveSeekerNet reveals the host-adaptation mechanisms at the codon level in IAV

The previous analysis demonstrated significant biases in nucleotide composition between avian and mammalian hosts. These biases appeared to be driven by differences in A/T and G/C content between mammalian-adapted and avian-adapted strains. Based on these results, we next examined codon-level signals associated with IAV host adaptation by computing SHAP values for ground-truth labels, revealing how the model “perceives” each host’s identity.

We began by assessing the relationship between codon usage bias and codon-level model importance. Fig. 5 (for 2 typical segments, PB2 and HA) and Additional file 1: Supplementary Fig. S8 (for the other 6 segments, PB1, PA, NP, NA, M, and NS) show that most codons exhibited highly significant associations between RSCU and SHAP values (q < 0.05). The left side of the volcano plot indicates a negative RSCU–SHAP association, suggesting that codons with higher usage tend to receive lower (more negative) SHAP values. A/T-rich codons were predominantly found on the negative correlation side for avian hosts, whereas G/C-rich codons increasingly occupied this region for mammalian hosts, with highly significant q-values (q < 0.05). However, many codons showed weak Spearman correlations (|𝜌| < 0.2), indicating only modest monotonic relationships between RSCU and model SHAP importance. Although individual Spearman correlations between RCSU and SHAP values were modest, their consistently negative direction across multiple codons suggests a systematic host signal with a cumulative effect. In SHAP analysis, these codons collectively contribute negative importance, suggesting that the model penalized A/T composition in avian hosts because higher A/T content is characteristic of mammalian hosts, and penalized G/C composition in mammalian hosts because higher G/C content is characteristic of avian hosts.

**Figure 5.**
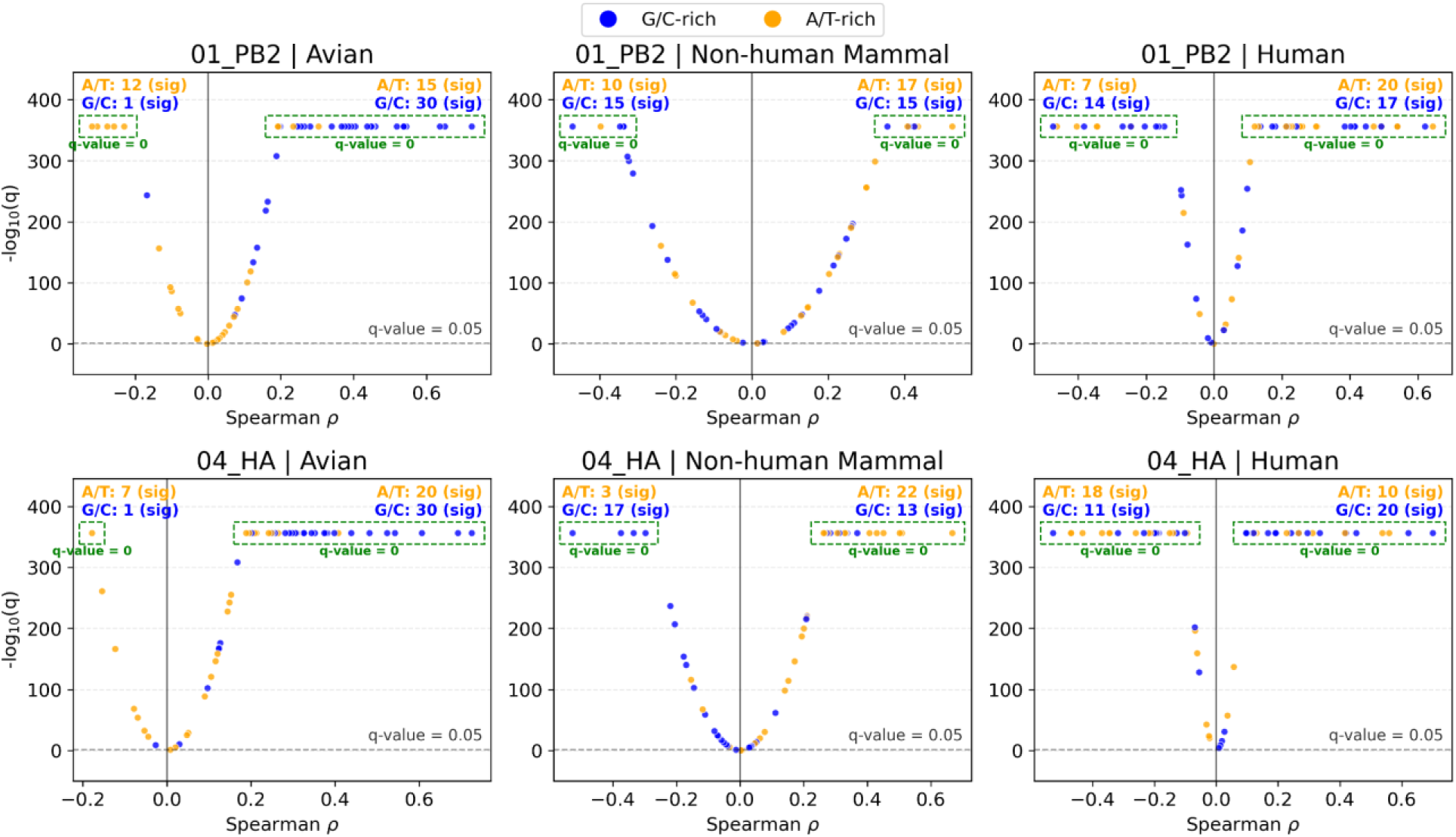
Volcano plot of Spearman correlation between RCSU and codon-SHAP values for segments PB2 and HA. Significance is expressed as − log_10_(𝑞). In each panel, the top-left and top-right annotations report the numbers of significant (q < 0.05) A/T-rich and G/C-rich codons on the negative- and positive-correlation sides, respectively. Blue and orange points indicate G/C-rich and A/T-rich codons, respectively.

On the other hand, the right side of the volcano plots indicates a positive RSCU–SHAP association, in which codon usage increases tend to receive higher (more positive) SHAP values. The “right-host” composition—predominant G/C-rich codons for avian hosts and increasing A/T-rich codons for mammalian hosts—resided in this region, with highly significant q-values (q < 0.05). Compared to the left side of the volcano plot, many codons showed stronger Spearman correlations (0.2 < 𝜌 < 0.8). In this context, the model assigned higher (positive) SHAP values to G/C-rich codons in avian hosts, reflecting their enrichment in the sequence and their role in driving avian classifications, whereas mammalian hosts are assigned higher (positive) SHAP values to A/T-rich codons, consistent with their more A/T-biased codon usage and their contribution to mammalian predictions.

To support the results above, we generated trajectory plots showing how SHAP value changes varied across host groups (Fig. 6). These patterns align with the volcano plot interpretation: the model systematically penalizes “wrong-host” composition (left side) while rewarding “right-host” composition (right side). Many A/T-rich lines (orange) were at or below zero in avian hosts and then increased in strains isolated from mammalian hosts, indicating that these codons contributed less (or negatively) to avian predictions but gained importance in mammals. By contrast, many G/C-rich lines (blue) started positive in avian hosts and then declined in mammalian hosts, indicating that G/C-rich codons were rewarded with higher SHAP values in the avian hosts but became less supportive of mammalian hosts. Because of these trajectories, many A/T-rich lines often rose and remained positive in mammalian hosts, whereas many G/C-rich lines crossed downward to zero or drifted negative, consistent with A/T-rich codons being rewarded and G/C-rich codons being penalized in mammalian predictions.

**Figure 6.**
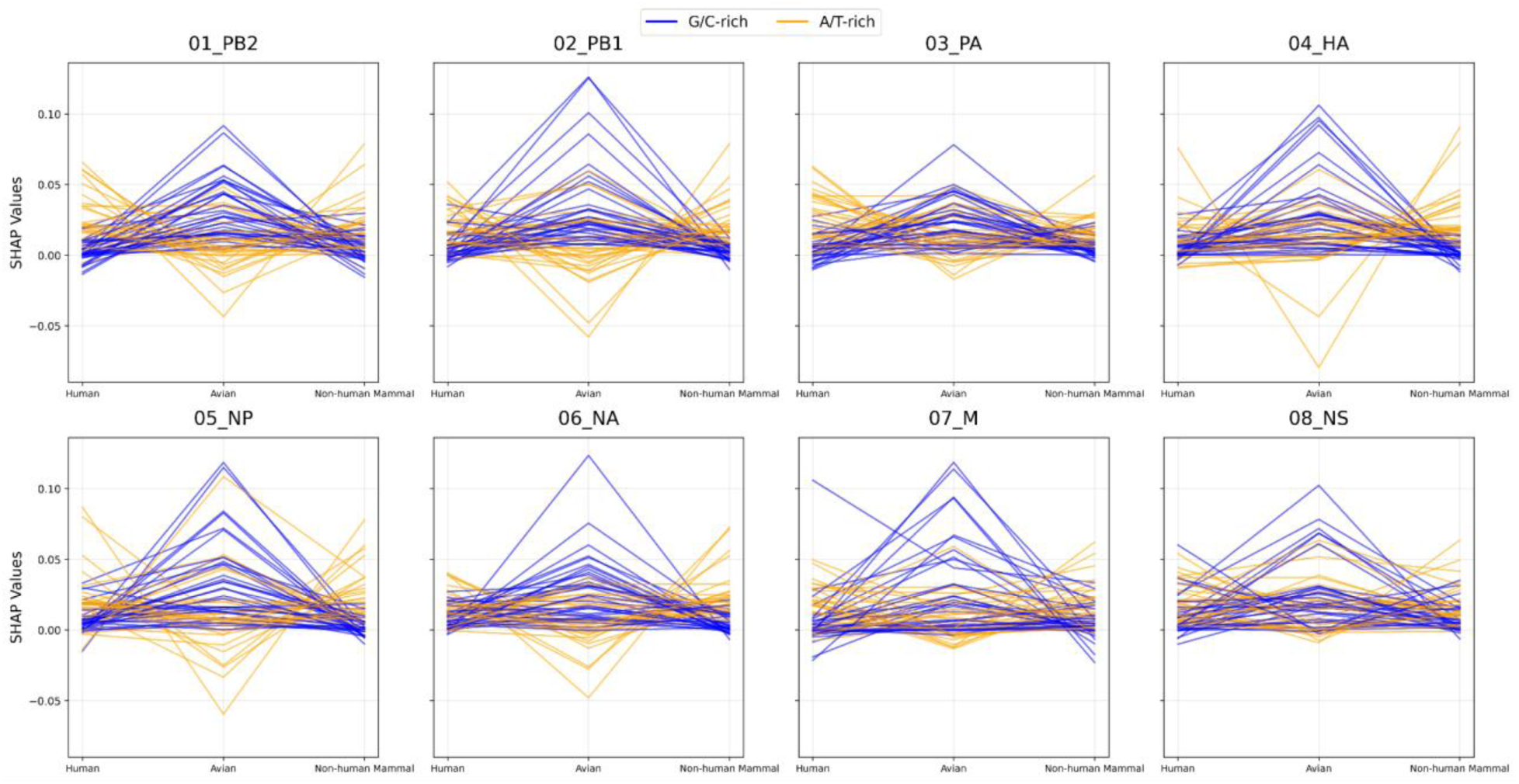
Trajectory plots of codon-SHAP value changes as the virus adapts from avian to mammalian hosts. Blue and orange lines denote G/C-rich and A/T-rich codons, respectively.

Avian and mammalian hosts exhibited distinct evolutionary trajectories. We next analyzed subtype-specific trajectories of codon-SHAP values in non-human mammalian hosts relative to human hosts, including H1N1, H3N2, H1N2, H2N2, H3N8, H7N7, H9N2, H7N9, H5N1, H5N6, and North American H5Nx subtypes (Additional file 1: Supplementary Figs. S9, S10, S11, and Fig.7). In avian-origin viruses (H5N1, H5N6, H9N2, H7N9, and North American H5Nx), many A/T-rich codons showed a downward trend, whereas many G/C-rich codons exhibited an upward trend; this convergence attenuated the divergence in SHAP values between A/T-rich and G/C-rich codons in human hosts (Fig. 7). The downward trend of A/T-rich codons was also observed in H1N1, H3N2, H1N2, H2N2, H3N8, and H7N7 (Additional file 1: Supplementary Figs. S9, S10, and S11). In contrast, many G/C-rich codons remained relatively stable, and some showed a slight upward trend in H1N1, H3N2, H1N2, and H2N2 (Additional file 1: Supplementary Figs. S9, S10), whereas they continued to decline in H3N8 and H7N7 (Additional file 1: Supplementary Fig. S11). Human hosts thus exhibited an attenuated divergence in codon-SHAP values between A/T-rich and G/C-rich codons, suggesting selective retention of certain G/C-rich codons alongside a decrease in A/T-rich favoritism. Nevertheless, A/T-rich codons displayed higher positive SHAP values than G/C-rich codons, supporting stronger associations with human-adaptive predictions.

**Figure 7.**
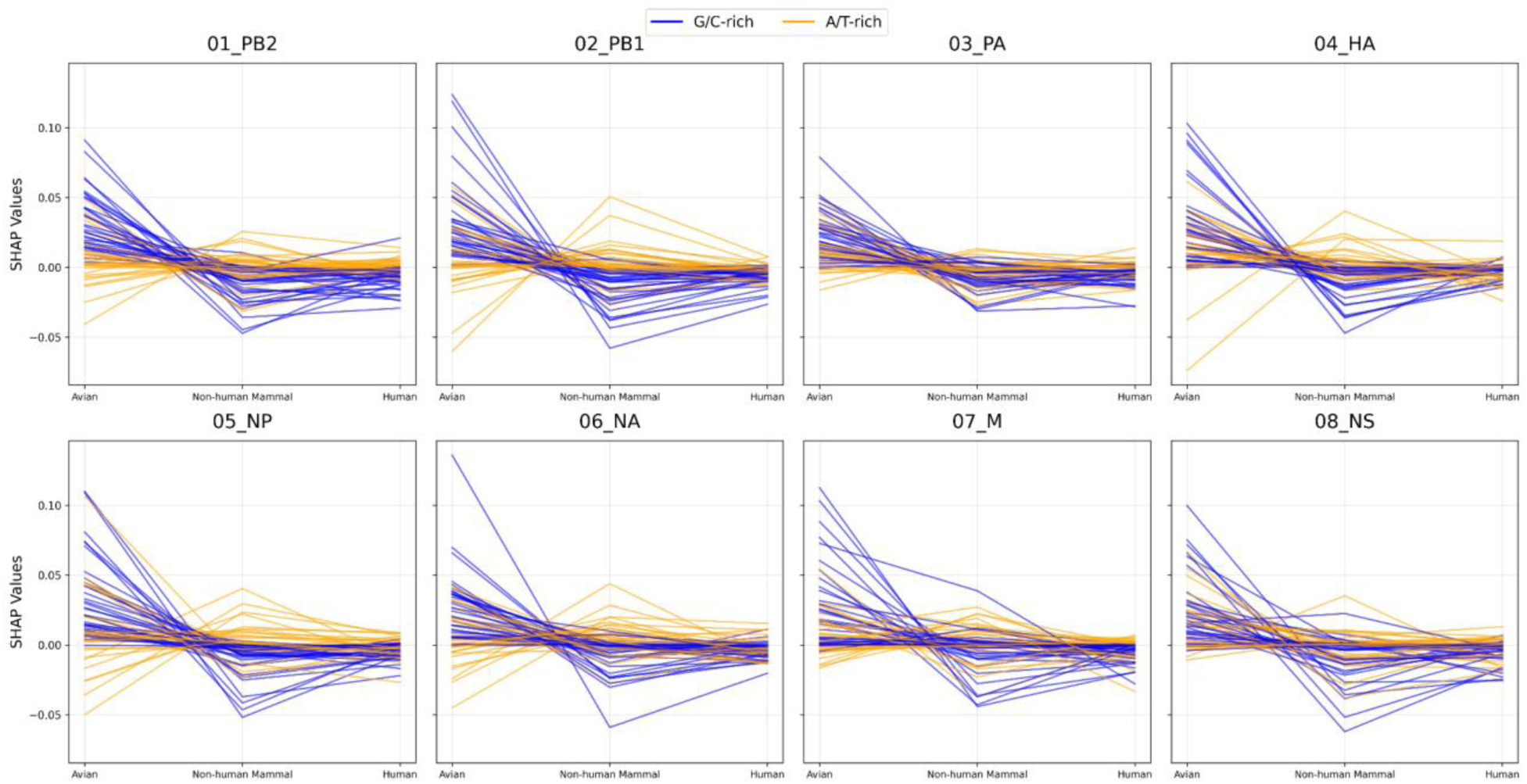
Subtype-specific (H5N1, H5N6, H9N2, H7N9, and North American H5Nx) trajectory plots. The plots illustrate changes of codon-SHAP values in non-human mammalian hosts relative to human hosts. Blue and orange lines denote G/C-rich and A/T-rich codons, respectively.

**Figure 8.**
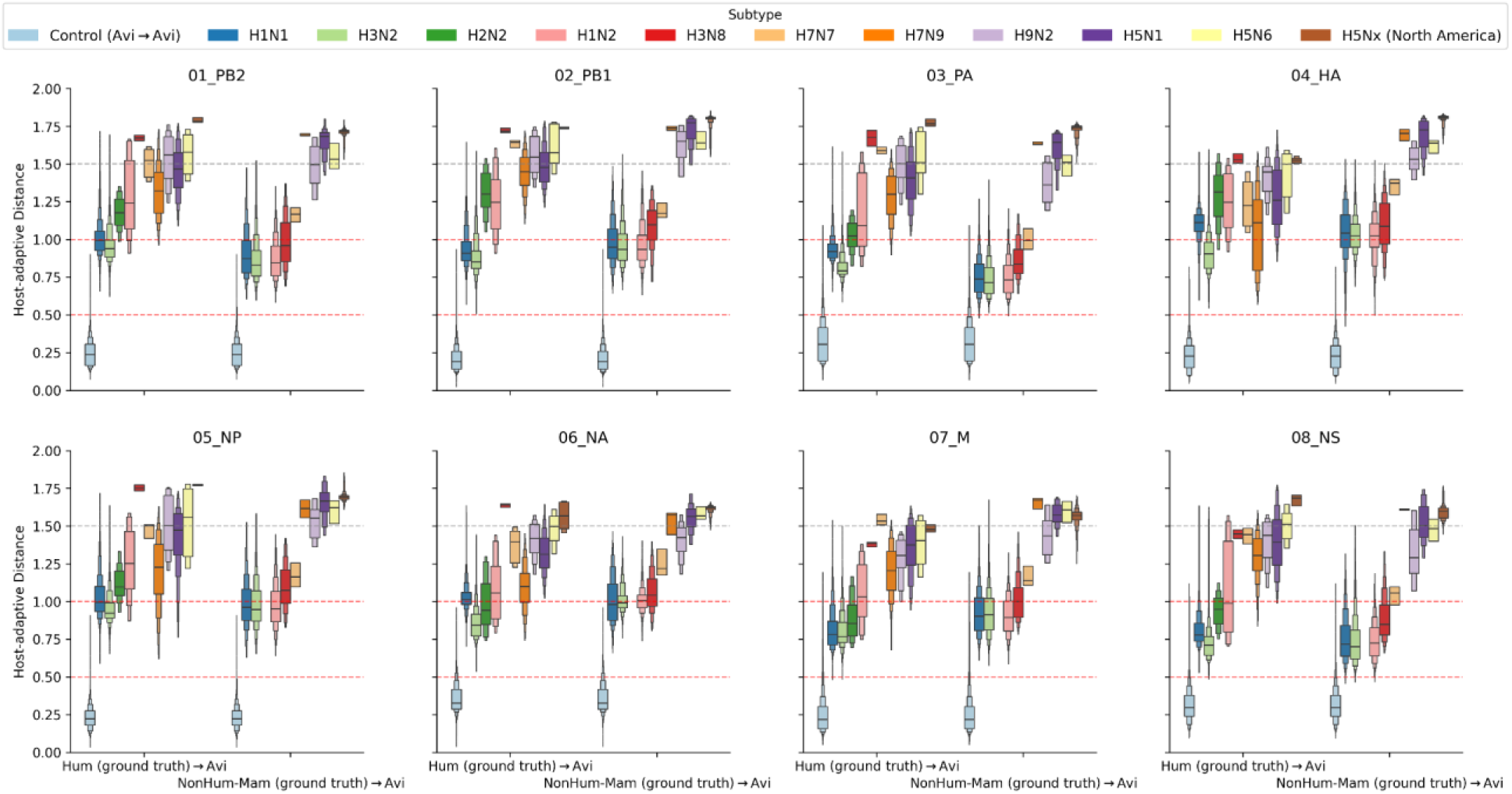
Host-adaptive distances relative to the global avian medoid using ground-truth codon-SHAP profiles. The boxen plots show human-to-avian, non-human mammal-to-avian, and avian-to-avian (control) host-adaptive distances. The global avian medoid was derived from the codon-SHAP profiles of pre-2020 avian strains. Red lines demarcate the “Mammalian Adaptation Zone,” while the distance between the upper red line and the gray line represents the hypothesized “Hard Distance.” The first set of boxen plots (left side) shows human-to-avian host-adaptive distances, and the second set (right side) shows non-human mammal-to-avian host-adaptive distances.

**Figure 9.**
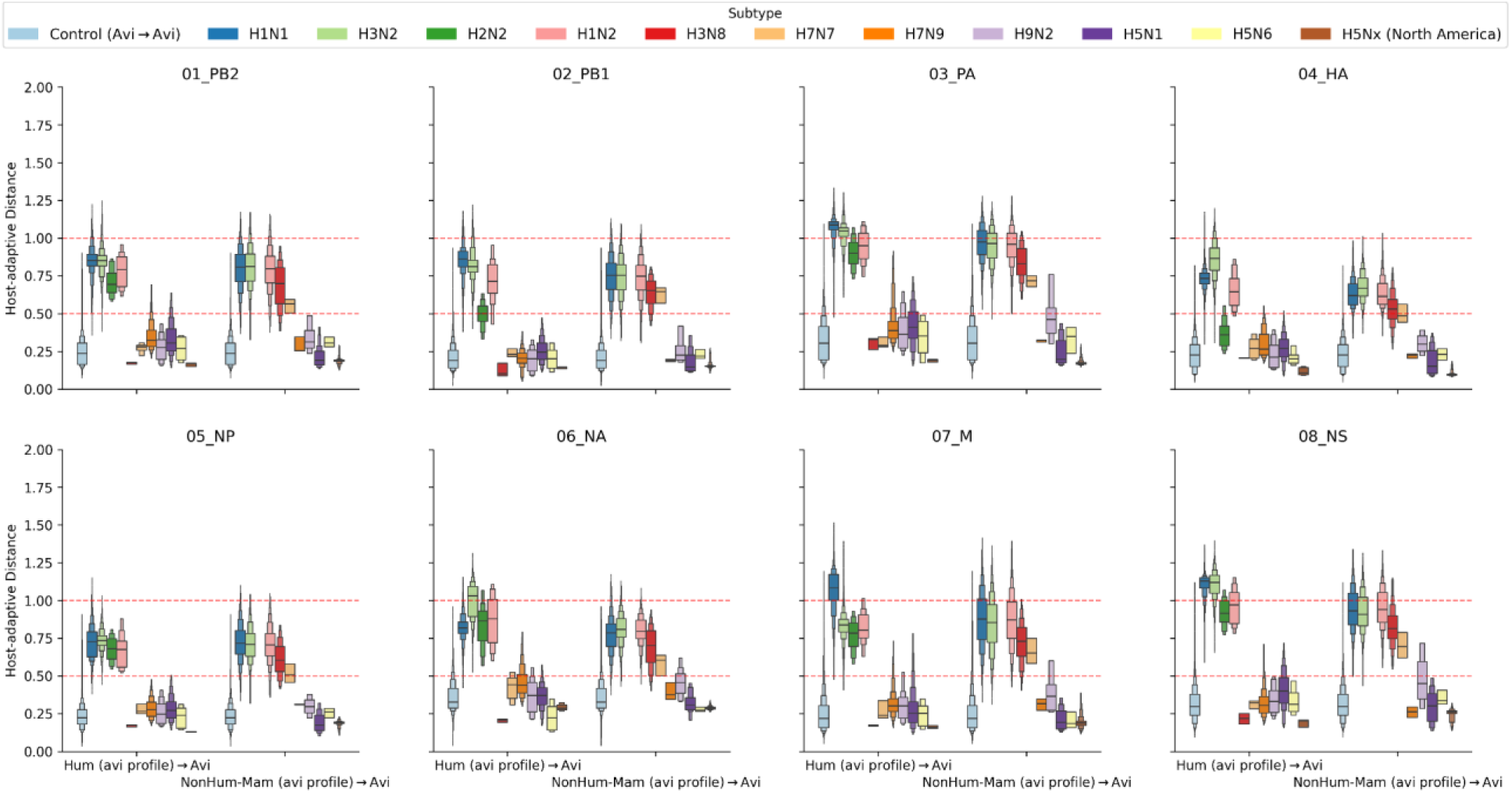
Host-adaptive distances relative to the global avian medoid using avian codon-SHAP profiles. The boxen plots show human-to-avian, non-human mammal-to-avian, and avian-to-avian (control) host-adaptive distances. The global avian medoid was derived from the codon-SHAP profiles of pre-2020 avian strains. Red lines demarcate the “Mammalian Adaptation Zone.” The first set of boxen plots (left side) shows human-to-avian host-adaptive distances, and the second set (right side) shows non-human mammal-to-avian host-adaptive distances.

**Figure 10.**
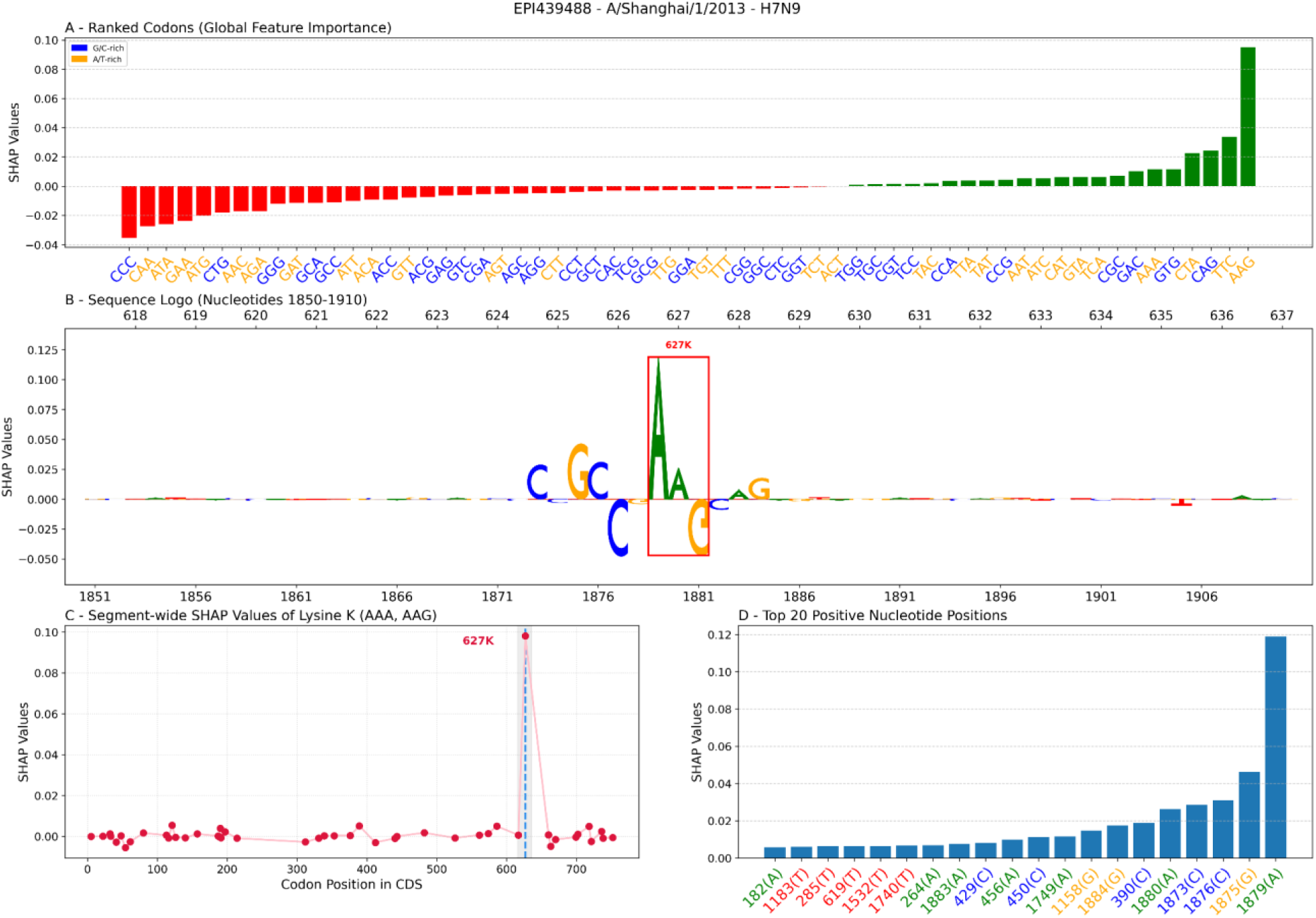
SHAP analysis of human isolate *A/Shanghai/1/2013* (EPI439488) showing WaveSeekerNet identifies PB2-E627K. Panel A shows the global feature importance (SHAP values) for 61 sense codons, ranked from the most negative to the most positive contribution to the human class. Panel B shows the SHAP values for each nucleotide of the AAG codon as a sequence logo of model importance scores. Panel C shows the segment-wide SHAP values of lysine (K), encoded by AAA or AAG, with a clear peak at position 627. Panel D shows the top 20 positively contributing nucleotide positions.

Taken together, these results demonstrate that codons characteristic of a given host (G/C-rich in avian hosts, A/T-rich in mammalian hosts) tend to shift toward positive SHAP in that host and toward negative SHAP in the others. Additionally, their selective retention in human hosts directly illustrates IAV’s host-adaptation mechanisms.

### 3.5 Quantifying host-adaptive distance in IAV

In this analysis, we apply our host-adaptive distance metric to track transitions in IAV genomic signatures across subtypes within host-adaptive space, hypothesizing a quantifiable barrier that the virus must breach to establish a persistent mammalian lineage. We established the avian-ancestor baseline using the codon-SHAP profiles of pre-2020 avian strains. This baseline represents the reservoir’s consensus evolutionary strategy—the foundational state from which zoonotic strains diverge. To characterize this state, we computed the global avian medoid and subtype-specific medoids, providing both a general and a lineage-specific reference for measuring adaptive evolution. Unlike a mean centroid, the medoid is an actual biological sequence (a representative member), ensuring our reference reflects a viable, functionally robust viral phenotype rather than a theoretical average. We then quantified host-adaptive distance as the cosine distance (Equation 5) between the codon-SHAP profiles of each query strain (e.g., human or non-human mammal queries) and the avian medoid. This metric captures divergence in the functional feature space, effectively measuring how far a virus has evolved from its ancestral avian identity. The analyses included H1N1, H3N2, H2N2, H1N2, H3N8, H7N7, H7N9, H9N2, H5N1, H5N6, and North American H5Nx subtypes.

We first calculated avian-to-avian (control) host-adaptive distances for all avian strains in our dataset, thereby defining the avian zone within host-adaptive space. Avian strains clustered within a host-adaptive distance range of 0.0–0.5 (the avian zone), which lies below the lower red line in Fig. 8 and clearly distinguishes avian host-adaptive space from mammalian host-adaptive space. This also reflected the model’s reliability (absolute MCS ≤ 0.02; Additional file 3: Supplementary Table S2) in distinguishing avian from mammalian hosts. We next calculated host-adaptive distances for mammalian hosts using the codon-SHAP profiles derived from their ground-truth profiles (Fig. 8 and Additional file 1: Supplementary Fig. S12). Analyses of persistent mammalian lineages—including H1N1, H3N2, and H2N2—alongside persistent non-human mammalian lineages such as swine H1N2 and equine influenza viruses (H7N7 and H3N8)—revealed that these strains consistently shifted toward, and often clustered within, a host-adaptive distance zone of 0.5–1.0 (between the two red lines). We hypothesize that this zone represents the “Mammalian Adaptation Zone” (MAZ), where viruses shifted their host-adaptive space to reach and establish persistent mammalian lineages. For example, the segment PB2 of human H1N1 viruses exhibited a mean host-adaptive distance of 1.0089 ± 0.1313, with a 75th percentile of 1.0563 and a 95th percentile of 1.2415 (Additional file 4: Supplementary Table S3). Notably, human H7N9 viruses—historically associated with severe outbreaks with high mortality rate in China in 2013 [38]—also tended to cluster within this range, distinguishing them from avian-origin counterparts such as H5Nx and H9N2. More specifically, we identified 25 PB2 sequences from this outbreak carrying the PB2-E627K mutation and found that their host-adaptive distances (Human (ground truth) → Avian) showed a tendency to approach 1.0, although they did not fully converge (mean = 1.1924 ± 0.1580, ranging from 0.8892 to 1.4882; see Additional file 5: Supplementary Table S4 for details of the host-adaptive distances of these sequences).

In contrast, mammalian isolates of avian-origin viruses (e.g., H9N2, H5N1, H5N6, and the North American H5Nx) exhibited substantial host-adaptive distances, with values predominantly exceeding 1.5 and approaching the observed upper bound (∼1.8). For example, the segment PB2 of non-human mammalian North American H5Nx viruses exhibited a mean host-adaptive distance of 1.7107 ± 0.0319, with a 75th percentile of 1.7234 and a 95th percentile of 1.7402 (Additional file 4: Supplementary Table S3). This degree of divergence indicated near-opposing trajectories between avian and mammalian host-adaptive spaces in avian-origin viruses. Their mammalian signatures have not yet transitioned toward the MAZ; instead, these viruses remain confined within the avian zone. This interpretation was consolidated by our model predictions, which show that avian-origin viruses isolated from mammals were classified as avian in nearly 100% of cases (Additional file 1: Supplementary Fig. S13). To further support this point, we specially analyzed 74 North American non-human mammalian PB2 sequences (e.g., dairy cow, harbor seal) that had lysine (K) at position 627, encoded by the AAA codon, and found that their host-adaptive distances (Non-human Mammal (ground truth) → Avian) were far from 1.0 (mean = 1.7230 ± 0.0363; see Additional file 5: Supplementary Table S4 for details of the host-adaptive distances of these sequences). The large discrepancy—a gap of > 0.5 between avian-origin spillovers (e.g., H9N2, H5Nx) and the successful establishment cluster (e.g., H1N1, H3N2)—constitutes the hypothesis of “Hard Distance,” which may prevent these viruses from establishing persistent mammalian lineages. This hypothesis was further consolidated by analyzing human spillover cases of other subtypes, specifically H7N7 and H3N8. Although H7N7 and H3N8 were enzootic in equines, the rare human isolates of these subtypes exhibited Host-adaptive distances exceeding 1.5 from the avian baseline, and the model identified all of them as avian origin, suggesting that these specific spillover strains retained a maladaptive, non-human phenotype and failed to converge toward the MAZ.

To consolidate the hypotheses of the MAZ and the “Hard Distance,” we subsequently calculated host-adaptive distances using codon-SHAP profiles derived from the avian class (avian profiles) instead of from the ground-truth labels. As illustrated in Fig. 9 and Additional file 1: Supplementary Fig. S14, the host-adaptive distances of persistent mammalian lineages (H1N1, H3N2, H1N2, H2N2, H7N7/equine, and H3N8/equine) were consistently clustered within the MAZ, suggesting that avian signatures of these viruses have escaped the avian zone and established distinct mammalian host-adaptive states. We observed an exception in segment HA of human H2N2 viruses, with most remaining confined within the avian zone. This aligned with the model’s predictions, which identified 77 of 100 human H2N2 sequences as avian origin, yet these viruses still trended toward the 0.5 threshold. In contrast, avian-origin viruses (e.g., H9N2, H7N9, and H5Nx) remained confined within the avian zone, below the lower red line. For example, we again analyzed 74 North American non-human mammalian PB2 sequences using their avian codon-SHAP profiles. Their host-adaptive distances (Non-human Mammal (avian profile) → Avian) were comparatively small (mean = 0.1723 ± 0.0178), indicating that these viruses retained avian signatures, and thus remained confined within the avian zone. Similarly, analyses of 25 PB2 sequences from the H7N9 human outbreak in China in 2013 revealed that their host-adaptive distances (Human (avian profile) → Avian) were smaller than 0.5 (mean = 0.3419 ± 0.0529), indicating that this virus was still confined within the avian zone despite having caused severe outbreaks in humans.

Taken together, persistent mammalian lineages converged toward a host-adaptive distance zone of 0.5–1.0 under both ground-truth and avian profiles. In contrast, unestablished mammalian lineages diverged from this range, failing to reach the upper bound of 1.0 (“Hard Distance”) with their own profiles and remaining below the lower bound of 0.5, within the avian zone, when evaluated with avian profiles. The observed convergence across multiple subtypes within a host-adaptive distance range of 0.5–1.0 (MAZ) suggests that this zone within host-adaptive space represents a general mammalian competency threshold—a quantifiable boundary that viruses typically cross when successfully breaching the species barrier to establish sustained lineages or long-term spillovers in non-avian hosts.

### 3.6 WaveSeekerNet identifies known changes impacting host adaptation

In this analysis, we demonstrate WaveSeekerNet’s ability to identify known changes that impact host adaptation. One well-known mutation that facilitates adaptation to mammalian hosts is the E627K mutation in the polymerase basic protein 2 (PB2). This substitution, in which glutamic acid (E) is replaced by lysine (K) at position 627, contributes to the high polymerase activity and enhances the virus’s ability to replicate efficiently in mammalian hosts [38,39]. Glutamic acid (E) is encoded by the codons GAA or GAG, while lysine (K) is encoded by the codons AAA or AAG. In 2013, avian influenza A(H7N9) emerged in China, and the first human cases were reported [38,39]. In this outbreak, the PB2-E627K mutation was associated with the higher case fatality rate. To validate WaveSeekerNet’s sensitivity to this known marker of mammalian adaptation, we curated 25 PB2 sequences from human isolates collected during the 2013–2014 H7N9 outbreak [38]. Fig. 10 presents the SHAP analysis for the representative isolate, *A/Shanghai/1/2013* (EPI439488), and Additional file 5: Supplementary Table S4 details the results for the full cohort. In panel A, the AAG codon is identified as the highest positive contributor to the human class. This is corroborated by the segment-wide view in panel C, which shows that the SHAP values exhibit a dominant peak specifically at position 627 of the codon AAG. A granular analysis of nucleotide contributions (panels B and D) reveals that the model assigned the highest positive weight to the adenine (A) at position 1879. This nucleotide corresponded precisely to the G→A transition that drives the E627K mutation (GAG→AAG). However, the SHAP value of the AAG codon was not universally positive across the cohort. For instance, in isolate *A/Anhui/03/2013* (EPI447637), the AAG codon was assigned a marginal negative SHAP value (−0.0125). A granular analysis of the top 20 positive nucleotide positions revealed that the model’s attention remained correctly focused: the adenine (A) at position 1879 retained the highest positive SHAP value (0.0890) (Additional file 5: Supplementary Table S4). This indicated that WaveSeekerNet prioritized the specific functional nucleotide substitution (G→A at the first codon position, driving GAG→AAG change) as the primary driver of host adaptation, even when broader codon-level or contextual features provided conflicting or dampened signals.

We further analyzed non-human mammalian isolates (e.g., dairy cow, skunk, harbor seal) from the ongoing North American H5Nx outbreak and found that their PB2 sequences carry lysine (K) at position 627, encoded by the AAA codon. Our SHAP analysis identified this specific AAA codon as a dominant feature for the non-human mammalian class. For example, in isolate *A/Harbor_Seal/QC/FAV-0836-4/2022* (EPI3053167), the AAA codon at position 627 and the adenine (A) at position 1879 were assigned the highest positive SHAP values (Additional file 1: Supplementary Fig. S15). In addition to 627K, we observed that other isolates in this outbreak carry 701N, which is encoded by the AAC codon. PB2-D701N, together with PB2-E627K, is recognized as one of the key markers of mammalian adaptation in PB2 [40,41]. Additional file 1: Supplementary Fig. S16 presents the SHAP analysis for the representative dairy cow isolate, *A/dairy cow/USA/002644-003/2025* (EPI4097884), where the AAC codon at position 701 and the adenine (A) at position 2101 were assigned the highest positive SHAP values. Full results for other isolates are detailed in Additional file 5: Supplementary Table S4. Not all isolates in this dataset exhibited these canonical high-confidence signatures. For example, in the isolate *A/harbor seal/Maine/22-020983-002/2022* (EPI2110498), the specific features associated with mammalian adaptation were absent or non-predictive, in which the AAA codon at position 627 contributed a negligible SHAP value (−0.0031), and the adenine (A) at position 1879 was assigned a marginal negative value (−0.0082) (Additional file 5: Supplementary Table S4). Nevertheless, A/T-rich codons predominantly contributed positive SHAP values, and the AAA and AAC codons had the top-ranked compositional scores.

## 4 Discussion

In this study, we leveraged our recently developed model, WaveSeekerNet [16], to dissect its internal learning processes, thereby gaining further insight into the evolution and biology of IAV. This approach enabled us to gain deeper insights into the mechanisms underlying IAV host adaptation from a computational perspective, highlighting the potential of modern deep learning as a powerful tool for deciphering viral evolution and informing pandemic prevention strategies and vaccine development.

Modern neural networks are known to produce miscalibrated confidence scores, in which predicted probabilities do not align with the actual correctness frequencies. This can create potentially misleading downstream decisions in surveillance and pandemic risk assessment. We therefore explicitly calibrated the WaveSeekerNet model’s predicted probabilities, achieving both markedly improved calibration and enhanced discriminative performance over an already high-performing model, demonstrating that more trustworthy confidence estimates can lead to better discrimination. This was clearly illustrated with human H1N1 seasonal viruses belonging to clades 6B.1A.5a.2a and 6B.1A.5a.2a.1, which have emerged and expanded since the 2021–2022 influenza season [42,43]. In this study, the WaveSeekerNet model was trained on pre-2020 sequences to predict post-2020 variants. Therefore, the absence or scarcity of sequences from these clades in the training set likely contributed to reduced accuracy in host-source prediction for clades 6B.1A.5a.2a and 6B.1A.5a.2a.1 human seasonal influenza cases. Furthermore, our analysis of host-adaptive distance revealed overlapping regions within the MAZ between the human-adapted and non-human mammalian-adapted zones (Figs. 8 and 9). This suggests that the model may struggle to distinguish fine-scale human adaptations within the broader MAZ due to shared genetic features across mammalian hosts. Class-wise calibration further indicated systematic biases, with underconfidence in the human class and overconfidence in the non-human mammalian class. Despite these problems, the calibrated model demonstrated robust generalization, accurately characterizing antigenically drifted A(H1N1)pdm09 variants not represented in the training data.

In our analysis of the correlation between RSCU values and codon-level SHAP values, we observed a statistically significant monotonic relationship. However, the strength of this association for most codons was weak to moderate, indicating that codon usage bias accounts for only a small fraction of the variance in codon-level SHAP values—that is, the effect is real but modest. These results suggest that the model is not simply memorizing individual codons; rather, it is detecting host-specific compositional signatures that arise from the aggregation of many weak but directionally consistent signals across the genome. In line with this view, one study found that many human-adapted viral genes exhibit directional changes in codon usage over time, including reduced overall G/C content and decreased use of G at the third codon position, consistent with host-driven selection on genome-wide composition [44].

SHAP analysis revealed positive contributions from G/C features in avian hosts and from A/T in mammalian hosts, demonstrating that the WaveSeekerNet model captured a fundamental evolution of IAV associated with host adaptation. This pattern is consistent with the empirical observations that avian-adapted viruses tend to maintain higher G/C content compared to human-adapted viruses, whose genomes rather evolved toward elevated A/T and lower G/C over time post-spillover, as quantified by log-odds classifiers (e.g., segments PB2, PB1, and PA) that distinguish hosts with near-100% accuracy even at the segment level [45]. A plausible interpretation is that there may be a connection between host body temperature and genomic G/C content. The avian G/C preference may reflect mutational equilibrium associated with replication of avian-adapted viruses in higher-temperature environments (∼40–42°C), such as in avian hosts, where G/C-rich sequences could contribute to a greater thermodynamic stability and secondary RNA structural integrity relative to A/T-rich sequences. A study by Yakovchuk et al. demonstrated that base-stacking interaction dominates the thermal stability of the DNA double helix [46].

Although IAV has a negative-sense, single-stranded RNA genome, its replication and packaging are influenced by conserved RNA secondary structures, including stem-loops and panhandle motifs, which operate within ribonucleoprotein complexes alongside essential protein-RNA interactions [47,48]. Within these regions, G/C may facilitate stronger base-stacking interactions with adjacent bases, potentially contributing to the stabilization of functional RNA stem-loops at the elevated temperatures typical of avian hosts. A G/C-rich genome could therefore potentially help oppose the thermal melting of these secondary structures and promote more efficient RNA folding and function. The G/C-temperature correlation was also established in prokaryotes, where G/C content positively correlates with optimal growth temperature [49]. However, this link remains underexplored and shows complications in IAV, particularly with PB1. While avian-origin PB1 enables viral replication at elevated temperatures, studies using chimeric genes and targeted mutagenesis show that this resistance is sufficiently driven by specific protein-level configurations, such as the PB1 amino acid substitutions G180E and S394P [50]. In vivo, a ∼2°C simulated fever protected mice against severe disease from a wild-type human IAV, but a virus engineered with these two specific avian-origin PB1 mutations overcame this thermal defense and caused severe disease [50].

Conversely, human-adapted viruses (∼33–37°C) were associated with C→T/G→A substitution biases, leading to reduced G/C content compared to avian viruses [45]. A recent study of 115,520 IAV whole genomes demonstrated that persistent mammalian lineages (e.g., H1N1, H3N2, H1N2, H2N2, equine H3N8) exhibit a consistent decline in G/C content compared to viruses that only infect birds [51]. Notably, this study also revealed that HPAI 2.3.4.4b H5 viruses found in mink, foxes, and humans share genomic signatures associated with long-term mammalian persistence (reduced G/C content) [51]. One key mechanism that could explain these findings is ZAP-mediated CpG sensing, in which Zinc-finger Antiviral Protein (ZAP) acts as a cellular sensor that detects and defends against invading RNA viruses by targeting CpG dinucleotides in their viral transcripts [52]. Because of this sensing mechanism, vertebrate-infecting RNA viruses have evolved to suppress CpGs in their genomes to avoid detection by ZAP [52]. Additional support for our observations comes from experimental results using avian ZAPs, which show that avian ZAPs can bind more promiscuously throughout the genomes of RNA viruses since they are less selective than mammalian ZAPs toward CpG-enriched RNA [53,54]. Another hypothesized driver involves APOBEC-like deaminases, in which human APOBEC enzymes—lacking functional avian orthologs—could deaminate cytidine to uridine on viral replication intermediates, potentially yielding C→T/G→A mutations during human adaptation [45,55]. However, studies have shown that APOBEC3 proteins probably do not significantly inhibit IAV replication [56,57], suggesting a minimal direct impact—though APOBEC knockdown or knockout could clarify its effects on nucleotide bias [55]. Experimental evidence also points to another cytoplasmic protein, KHNYN, which can restrict CpG-enriched viral RNA independently of ZAP [54]. Interestingly, KHNYN is found in mammals but is absent in avian species, likely emerging as a result of a gene-duplication event that happened after mammals and birds diverged [54]. Together with the cytoplasmic isoform of ZAP, the activity of these proteins provides biological support for our results showing that mammalian-adapted IAVs have a CpG-depleted genomic signature.

The HPAI H5N1 clade 2.3.4.4b viruses, including those detected in U.S. dairy cattle, retain avian-like genomic signatures and exhibit genetic and structural profiles characteristic of the Eurasian goose/Guangdong (Gs/GD/96) lineage [58–60]. They also exhibited predominant avian-type receptor (𝛼2,3 sialic acid) preference [61]. Thus, despite unprecedented spillover and the recently reported low mortality bovine-to-bovine transmission [59,60], these viruses remain best classified as avian influenza viruses. IAV can acquire mutations, such as PB2-E627K and PB2-D701N, that enhance replication in mammalian hosts [38–41]. These changes do not automatically mean that the virus will evolve the ability to sustain a highly pathogenic mammalian transmission lineage with pandemic potential, only that they are mutations of concern and that these lineages should be monitored closely, given the ability of IAV to rapidly evolve and reassort. Therefore, there is a critical need for robust approaches to anticipate which avian viruses are approaching this transition to persistent mammalian circulation. Our definition of host-adaptive distance could fundamentally advance our monitoring of IAV evolution and enhance our ability to predict when the virus will establish a persistent mammalian lineage. North American H5Nx viruses exemplify this: our analysis revealed that they remain stalled at the “Hard Distance,” far from the MAZ red line. Continuous host-adaptive distance tracking is a quantitative measure that tracks the evolutionary trajectory of IAV toward or away from this red line, thereby indicating the potential for a strain to establish a persistent mammalian lineage.

## 5 Conclusions

By interpreting the WaveSeekerNet model, we uncovered novel biological insights into IAV host adaptation and demonstrated the value of interpretable machine learning for biological discovery. Our findings showed that WaveSeekerNet extended beyond prediction to generate biologically meaningful hypotheses, offering a deeper understanding of IAV evolution and its adaptation to mammalian hosts. The model revealed fundamental mechanisms underlying host adaptation and identified key mutations, such as PB2-E627K and PB2-D701N. Beyond these insights, we introduced a host-adaptive distance metric to track viral evolution, a critical component of IAV surveillance. Importantly, this study established a well-calibrated predictive model that provided reliable confidence estimates—an essential feature for high-stakes applications such as IAV surveillance. At the Canadian Food Inspection Agency (CFIA)–National Centre for Foreign Animal Disease (NCFAD), which houses the World Organization for Animal Health reference laboratory for avian influenza, we have developed and routinely use the CFIA-NCFAD/nf-flu workflow (https://github.com/CFIA-NCFAD/nf-flu) for IAV analysis and surveillance [62]. We will integrate well-calibrated WaveSeekerNet and host-adaptive distance metric into CFIA-NCFAD/nf-flu to enhance Canadian and global IAV surveillance with a particular focus on risk assessment for H5Nx viruses currently circulating in North America.

## Supplementary Information

**Additional file 1**

**Supplementary Figures 1 to 16**

**Additional file 2**

**Supplementary Table S1. Data distribution of the study. Additional file 3**

**Supplementary Table S2. Class-wise calibration results for the pre-2020 and post-2020 datasets.** Calibration is evaluated using ECE, MCS, MCE, and ACE. Directionality: BALANCED (well-calibrated; absolute MCS ≤ 0.02), OVERCONF (overconfident; MCS > 0.02), UNDERCONF (underconfident; MCS < -0.02).

**Additional file 4**

**Supplementary Table S3. MAZ estimation parameters derived from empirical host-adaptive distance distributions of human-to-avian and non-human mammalian-to-avian hosts.** The tables report host-adaptive distances from ground-truth and avian profiles at the 75th and 95th percentiles, along with median and mean distances.

**Additional file 5**

**Supplementary Table S4. Reports of SHAP analysis to identify PB2-E627K and PB2-D701N.** The tables include sequences from human H7N9 outbreaks in China in 2013 and from North American H5Nx viruses. SHAP values for codons at positions 627 and 701 are reported, along with the top 20 positively associated positions and corresponding host-adaptive distances relative to the global avian medoid.

## Abbreviations

IAV: Influenza A Virus
MAZ: Mammalian Adaptation Zone
PB2: Polymerase Basic Protein 2
PB1: Polymerase Basic Protein 1
PA: Polymerase Acidic Protein
HA: Hemagglutinin
NP: Nucleoprotein
NA: Neuraminidase
M: Matrix Protein
NS: Nonstructural Protein
HPAI: Highly Pathogenic Avian Influenza
SHAP: SHapley Additive exPlanations
VADR: Viral Annotation DefineR
BA: Balanced Accuracy
MCC: Matthews Correlation Coefficient
ECE: Expected Calibration Error
MCE: Maximum Calibration Error
MCS: Miscalibration Score
ACE: Adaptive Calibration Error

## Data Availability

Data and code are available at https://github.com/nhhaidee/IAV_Evolution. The whole genomes of IAVs can be downloaded from EpiFlu GISAID (https://gisaid.org/) after creating an account and accepting the terms of use.

## Funding

H-H.N. and J.R. are supported by the Canadian Safety and Security Program (CSSP) grant CSSP-2023-CP-2620. “Pilot Pan-Canadian Surveillance for Mammalian Viral Pathogens Using Hematophagic Organisms and Environmental Samples”, awarded to D.L. and O.L.. H-H.N is also partially supported by a CHIR Team Grant: Building capacity in interdisciplinary research on Mpox (monkeypox) and other (re)emerging zoonotic threats to health titled “Mpox exposure and transmission at the human-animal interface; a One Health approach to viral ecology”, awarded to S.M. and O.L.. H-H.N. is also partially supported by the Natural Sciences and Engineering Research Council of Canada (NSERC), awarded to C.K.L. and the University of Manitoba

## Author’s Contributions

H-H.N. collected/curated data, trained the model, performed data analysis, and wrote the manuscript. J.R. reviewed/edited the manuscript, mentored, and provided support and feedback. O.L., C.K.L., and Y.B. supervised the study and reviewed/edited the manuscript. D.L. and S.M. reviewed/edited the manuscript. All authors contributed to finalizing the manuscript.

## Acknowledgements

We gratefully acknowledge all data contributors, i.e., the Authors and their Originating laboratories responsible for obtaining the specimens, and their Submitting laboratories for generating the genetic sequence and metadata and sharing via the GISAID Initiative, on which this research is based. The authors thank Dr. Tamiru Alkie, Dr. Anthony Signore, and Dr. Oksana Vernygora at the Canadian Food Inspection Agency (CFIA)–National Centre for Foreign Animal Disease (NCFAD) for valuable comments and feedback.

## Disclosure of use of AI-assisted tools

In preparing this manuscript, the authors used Perplexity Pro and Grammarly to improve readability, refine language and flow, and check for spelling and grammatical errors. Prompts such as “check and improve sentences” were used with Perplexity Pro. All suggestions generated by these tools were reviewed and edited by the authors to ensure accuracy and consistency with the study’s analyses and findings.

## Declarations

### Ethics approval and consent to participate

Not applicable

### Consent for publication

Not applicable

### Competing Interests

The authors have declared that no competing interests exist.

## Notes

### Competing Interest Statement

The authors have declared no competing interest.

### Summary of Updates

- There are two main parts of this paper: model calibration for reliable predictions and interpretable model for biological insights, so the title is updated to fully reflect the contributions of this study. - Add Conclusion section - Fixed wordings (e.g., Figure title, legend, abstract), update Figure 1 to fully illustrate the framework.

