## Additional file 1 for "Explainable and Calibrated AI for Decoding Host-Adaptive Changes in Influenza A Virus"

**Supplementary Figure S1. Reliability diagrams for the post-2020 dataset, constructed using 10 probability bins.** The diagonal line denotes perfect calibration, and the pink and blue curves show model calibration before and after calibration, respectively. Blue boxes highlight bins with large calibration gaps (> 0.05).

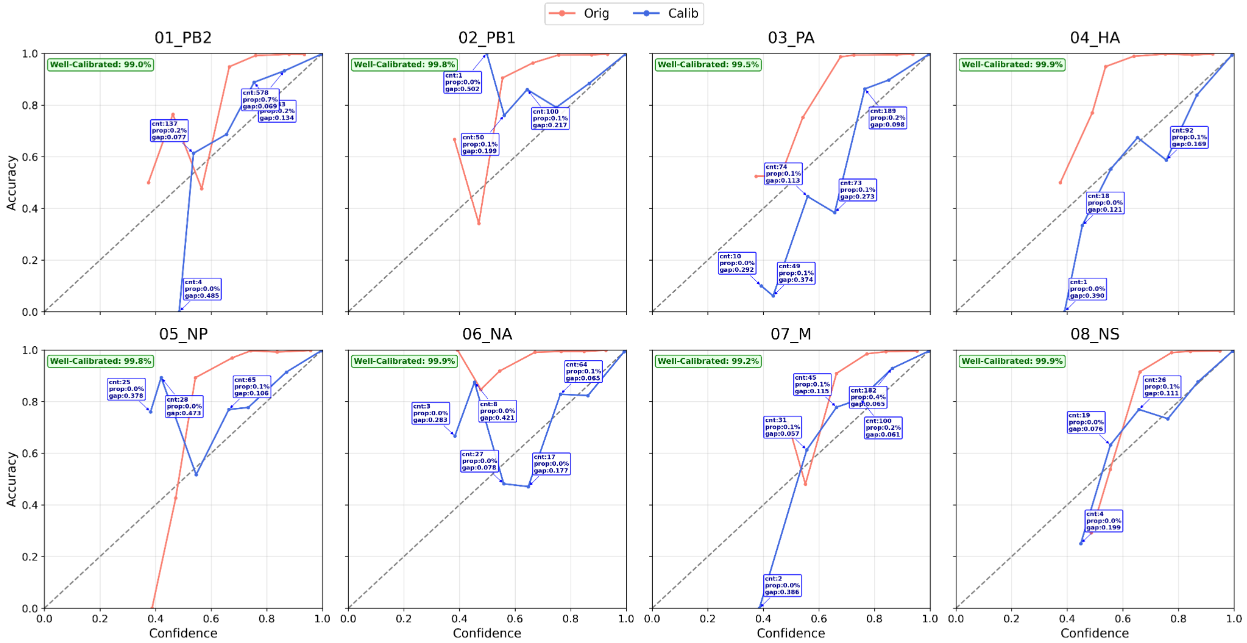


**Supplementary Figure S2. Model performance on the pre-2020 dataset.** Bar colors indicate: pink, before calibration; blue, after calibration. Performance is evaluated using F1-score (Macro Average), BA, and MCC.

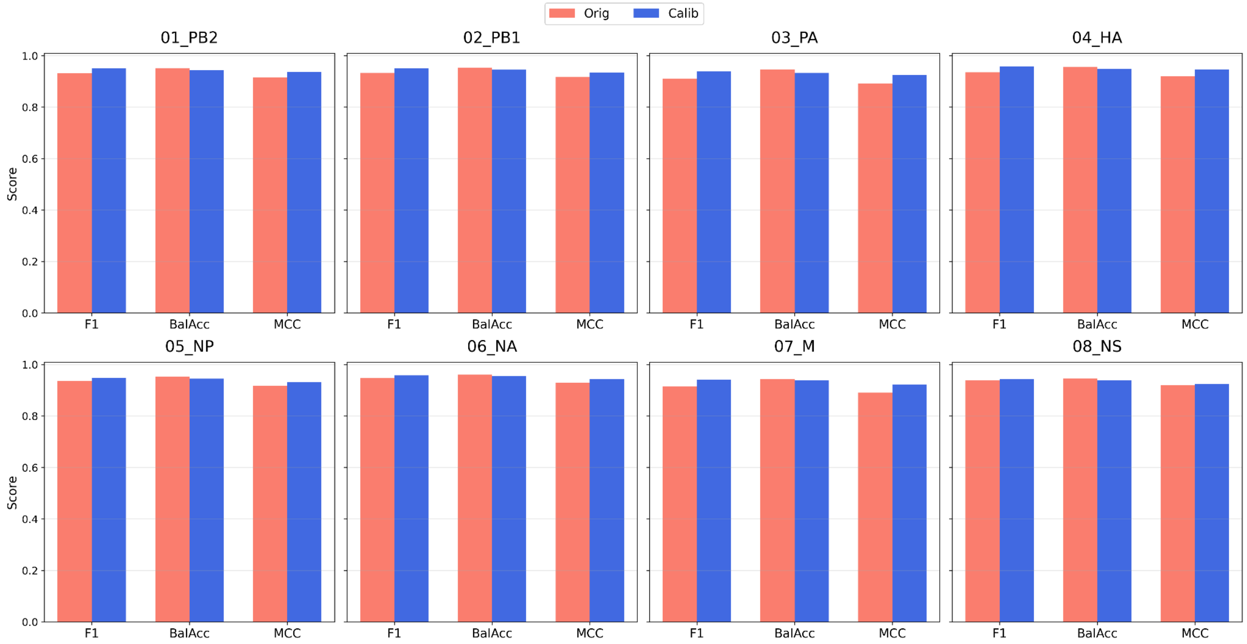

**Supplementary Figure S3: Calibration results for the pre-2020 dataset.** Calibration is evaluated using ECE, MCS, MCE, and ACE.

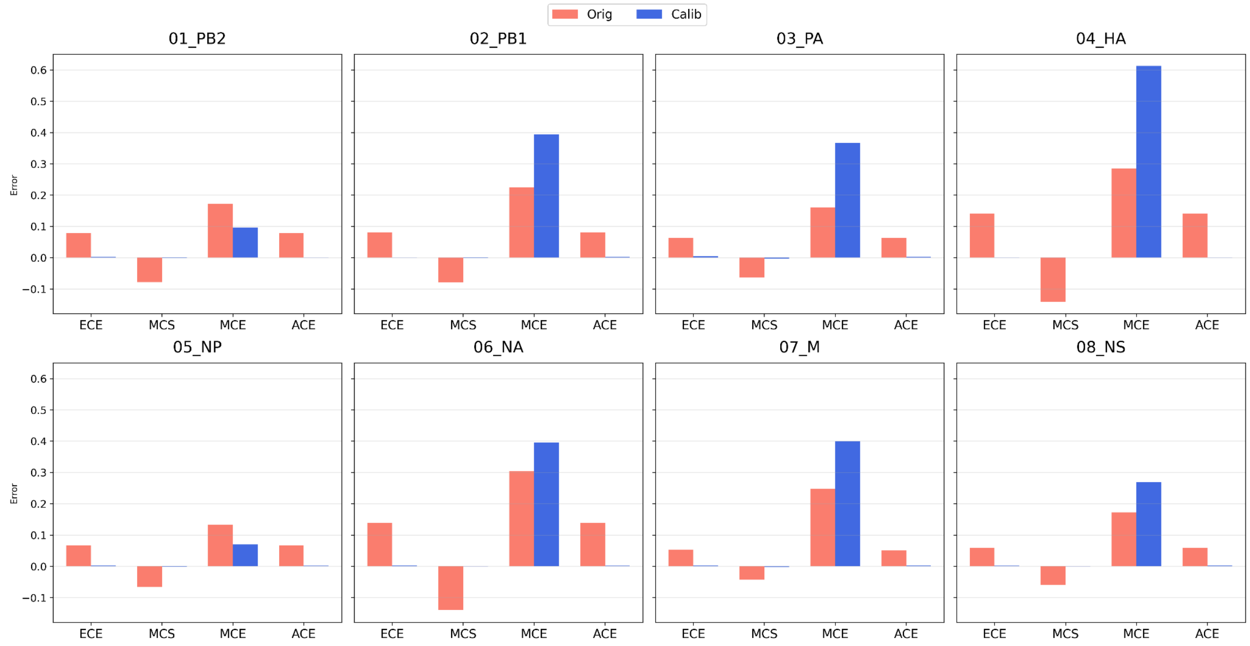


**Supplementary Figure S4. Reliability diagrams for the pre-2020 dataset, constructed using 10 probability bins.** The diagonal line denotes perfect calibration, and the pink and blue curves show model calibration before and after calibration, respectively. Blue boxes highlight bins with large calibration gaps (> 0.05).

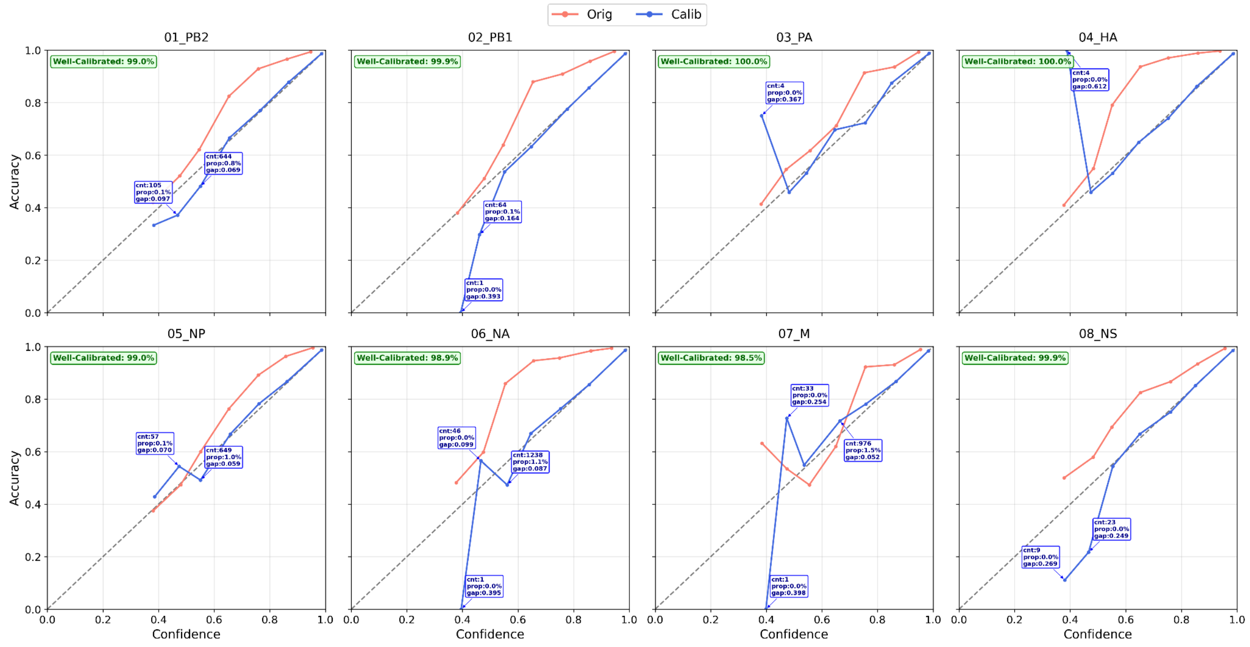

**Supplementary Figure S5: Distribution of nucleotide-SHAP values across host groups and genome segments on the pre-2020 dataset.** Boxen plots show SHAP value distributions for correctly predicted sequences.

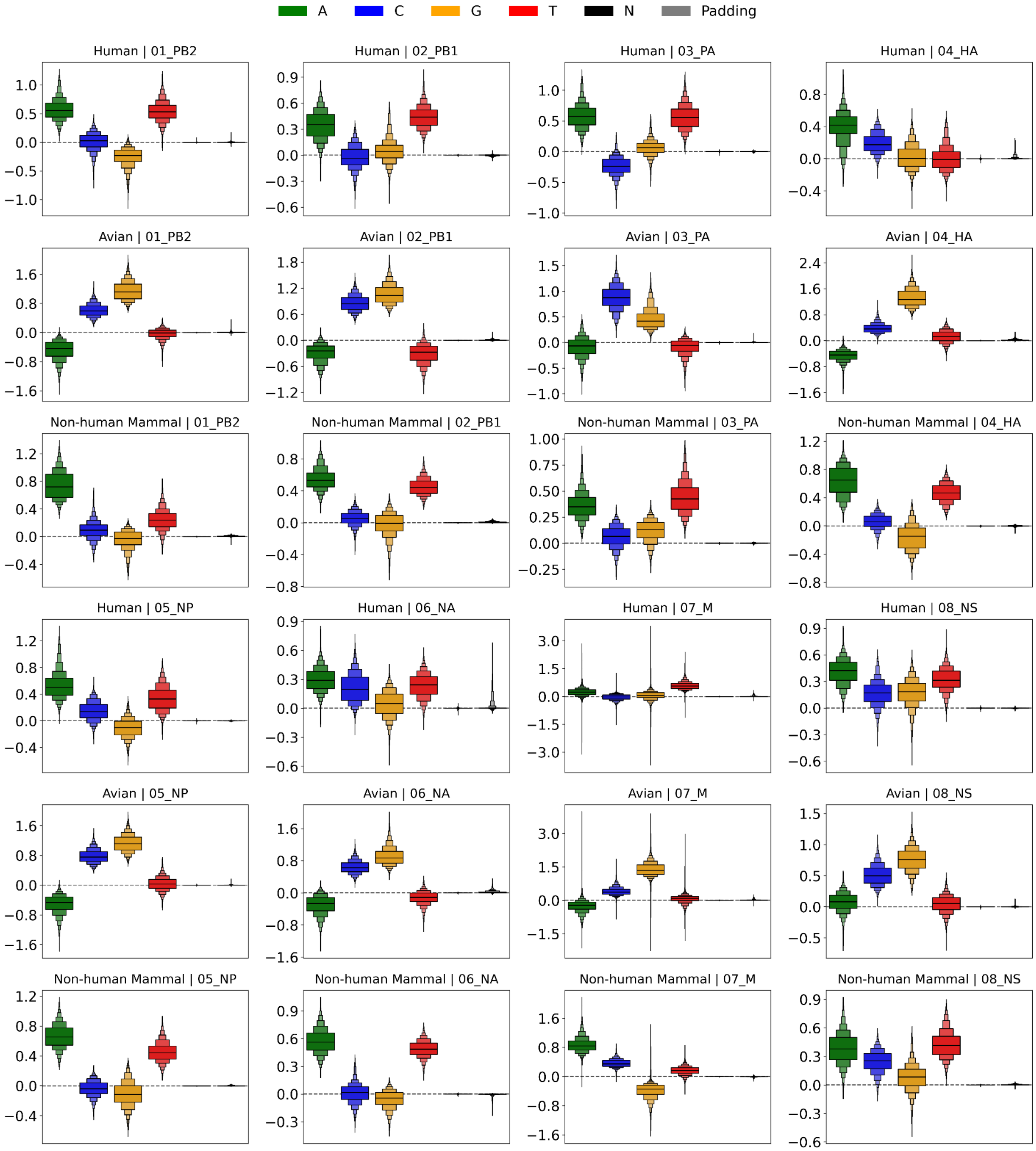

**Supplementary Figure S6. Distribution of nucleotide-SHAP values across host groups and genome segments on the pre-2020 dataset.** Boxen plots show SHAP value distributions for sequences with host-source discrepancies (mismatches between the predicted and known host sources).

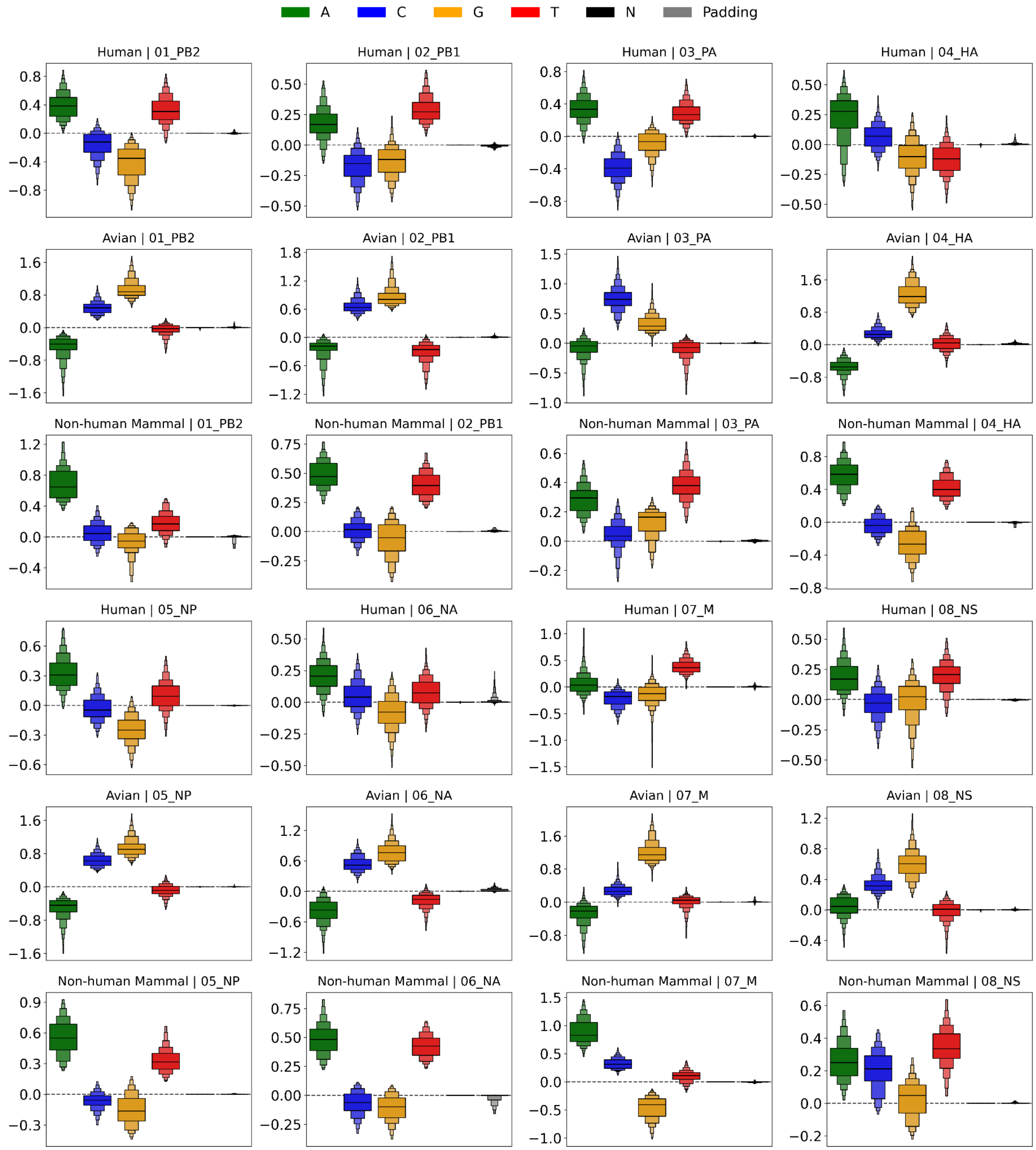

**Supplementary Figure S7: Distribution of nucleotide-SHAP values across host groups and genome segments on the post-2020 dataset.** Boxen plots show SHAP value distributions for sequences with host-source discrepancies (mismatches between the predicted and known host sources), aggregated across 10 cross-validation folds.

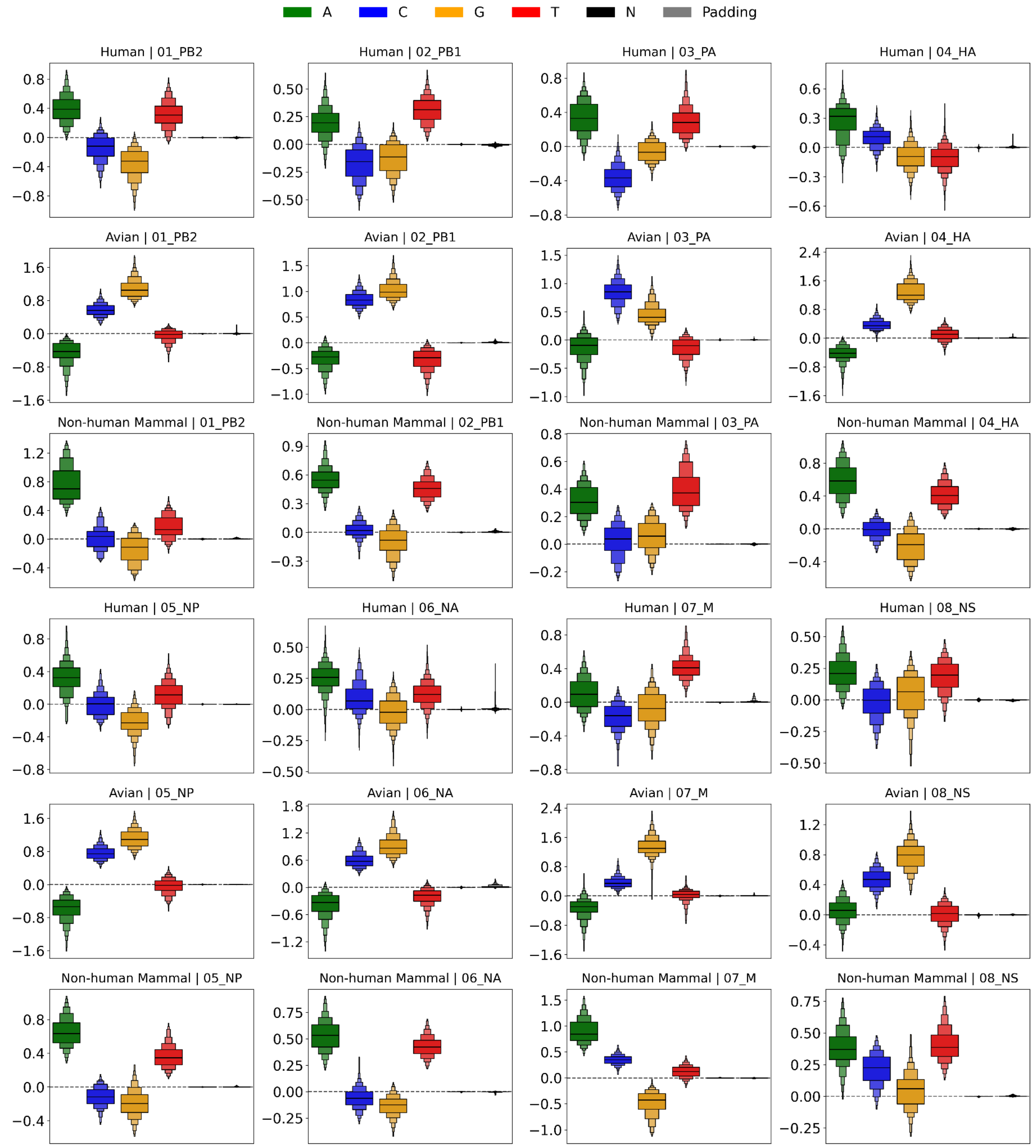

**Supplementary Figure S8. Volcano plot of Spearman rank correlation between RCSU and codon-SHAP values for segments PB1, PA, NP, NA, M, and MS**. Significance is expressed as $-\ log_{10}(q)$. In each panel, the top-left and top-right annotations report the numbers of significant ($q < 0.05$) A/T-rich and G/C-rich codons on the negative- and positive-correlation sides, respectively. Blue and orange points indicate G/C-rich and A/T-rich codons, respectively.

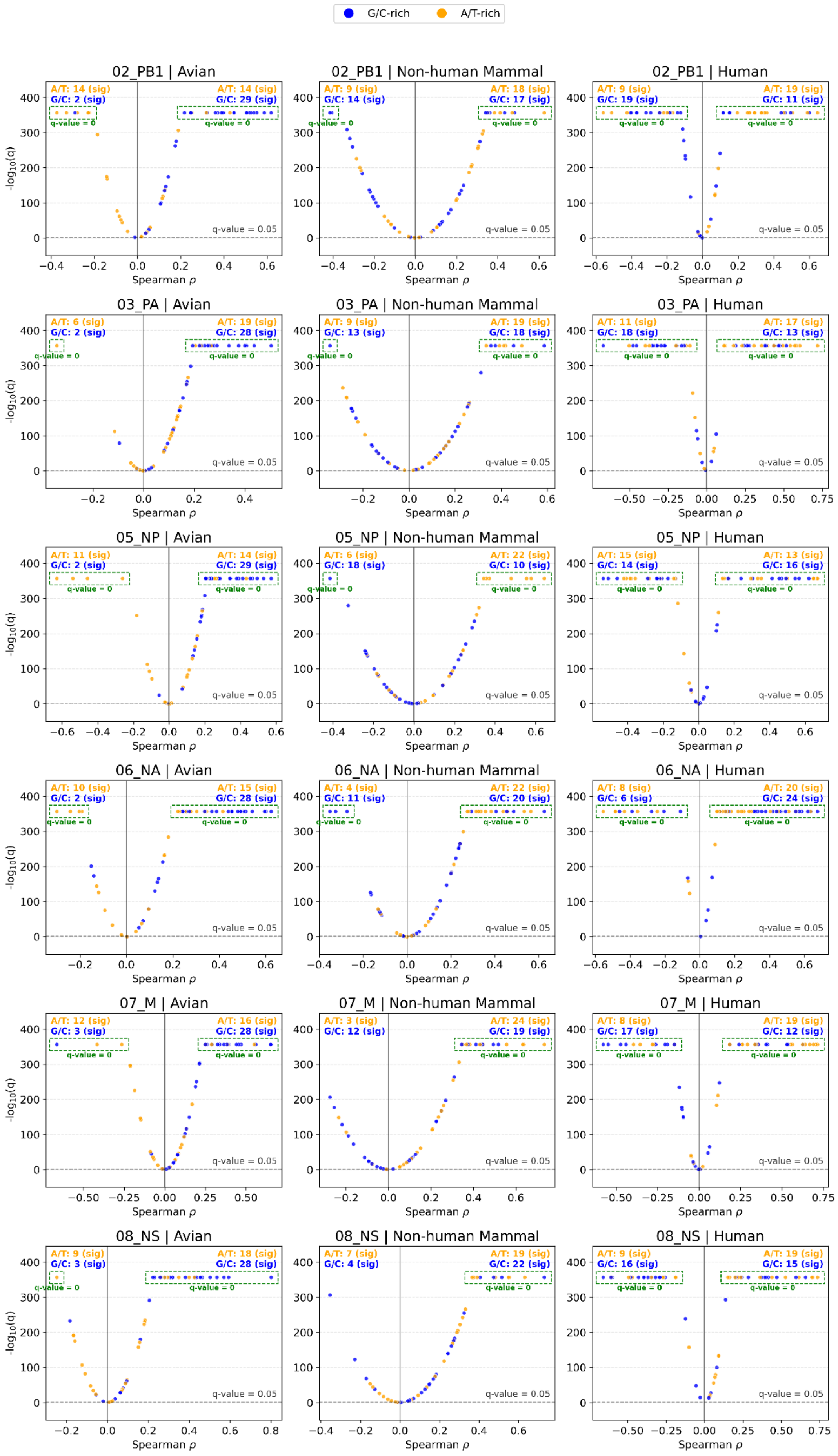
G/C-rich
A/T-rich
02_PB1 | Avian
A/T: 14 (sig)
G/C: 2 (sig)
A/T: 14 (sig)
G/C: 29 (sig)
02_PB1 | Non-human Mammal
A/T: 9 (sig)
G/C: 14 (sig)
A/T: 18 (sig)
G/C: 17 (sig)
02_PB1 | Human
A/T: 9 (sig)
G/C: 19 (sig)
A/T: 19 (sig)
G/C: 11 (sig)
03_PA | Avian
A/T: 6 (sig)
G/C: 2 (sig)
A/T: 19 (sig)
G/C: 28 (sig)
03_PA | Non-human Mammal
A/T: 9 (sig)
G/C: 13 (sig)
A/T: 19 (sig)
G/C: 18 (sig)
03_PA | Human
A/T: 11 (sig)
G/C: 18 (sig)
A/T: 17 (sig)
G/C: 13 (sig)
05_NP | Avian
A/T: 11 (sig)
G/C: 2 (sig)
A/T: 14 (sig)
G/C: 29 (sig)
05_NP | Non-human Mammal
A/T: 6 (sig)
G/C: 18 (sig)
A/T: 22 (sig)
G/C: 10 (sig)
05_NP | Human
A/T: 15 (sig)
G/C: 14 (sig)
A/T: 13 (sig)
G/C: 16 (sig)
06_NA | Avian
A/T: 10 (sig)
G/C: 2 (sig)
A/T: 15 (sig)
G/C: 28 (sig)
06_NA | Non-human Mammal
A/T: 4 (sig)
G/C: 11 (sig)
A/T: 22 (sig)
G/C: 20 (sig)
06_NA | Human
A/T: 8 (sig)
G/C: 6 (sig)
A/T: 20 (sig)
G/C: 24 (sig)
07_M | Avian
A/T: 12 (sig)
G/C: 3 (sig)
A/T: 16 (sig)
G/C: 28 (sig)
07_M | Non-human Mammal
A/T: 3 (sig)
G/C: 12 (sig)
A/T: 24 (sig)
G/C: 19 (sig)
07_M | Human
A/T: 8 (sig)
G/C: 17 (sig)
A/T: 19 (sig)
G/C: 12 (sig)
08_NS | Avian
A/T: 9 (sig)
G/C: 3 (sig)
A/T: 18 (sig)
G/C: 28 (sig)
08_NS | Non-human Mammal
A/T: 7 (sig)
G/C: 4 (sig)
A/T: 19 (sig)
G/C: 22 (sig)
08_NS | Human
A/T: 9 (sig)
G/C: 16 (sig)
A/T: 19 (sig)
G/C: 15 (sig)
q-value = 0
q-value = 0.05
-log10(q)
Spearman ρ

**Supplementary Figure S9. Subtype-specific (H1N1 and H3N2) trajectory plots.** The plots illustrate changes of codon-SHAP values in non-human mammalian hosts relative to human hosts. Blue and orange lines denote G/C-rich and A/T-rich codons, respectively.

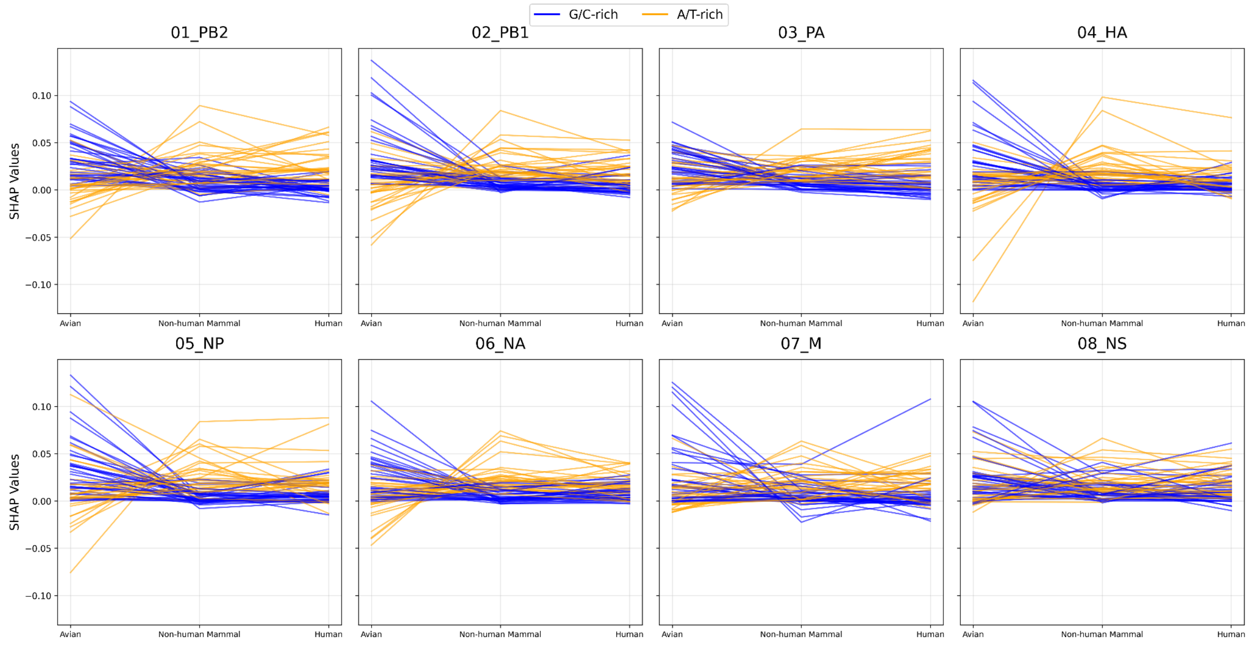


**Supplementary Figure S10. Subtype-specific (H1N2 and H2N2) trajectory plots.** The plots illustrate changes of codon-SHAP values in non-human mammalian hosts relative to human hosts. Blue and orange lines denote G/C-rich and A/T-rich codons, respectively.

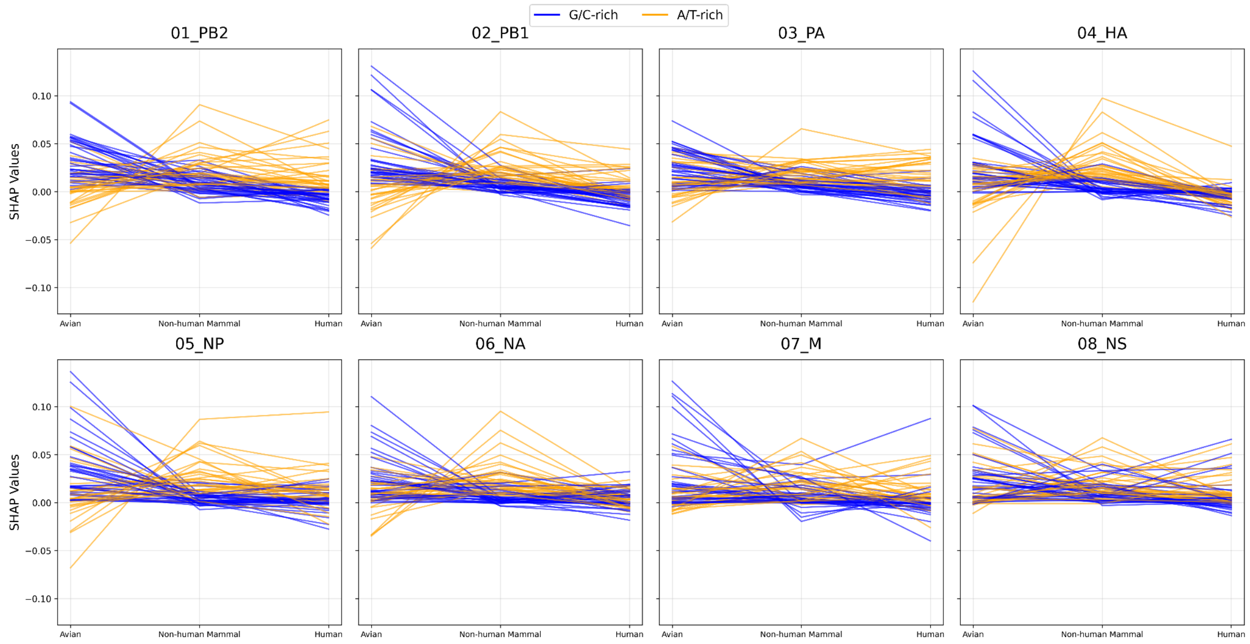

**Supplementary Figure S11. Subtype-specific (H3N8 and H7N7) trajectory plots.** The plots illustrate changes of codon-SHAP values in non-human mammalian hosts relative to human hosts. Blue and orange lines denote G/C-rich and A/T-rich codons, respectively.

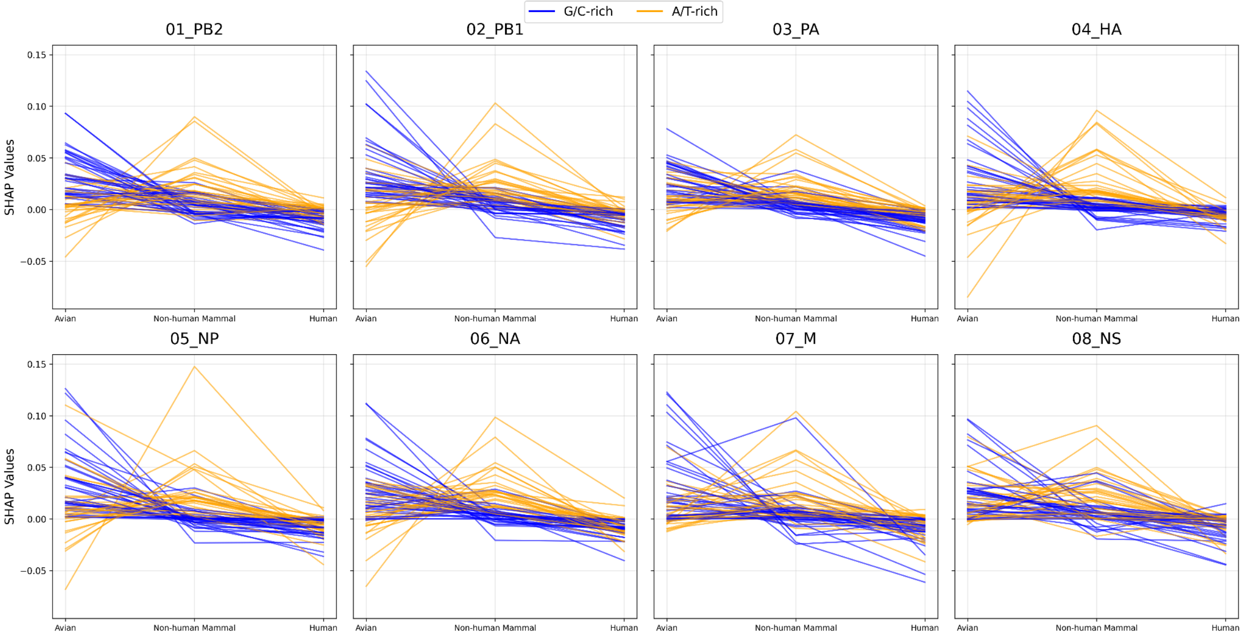

**Supplementary Figure S12. Host-adaptive distances relative to the subtype-specific medoids using ground-truth codon-SHAP profiles.** The boxen plots show human-to-avian, non-human mammal-to-avian, and avian-to-avian (control) host-adaptive distances. These medoids were derived from the codon-SHAP profiles of pre-2020 avian strains within each subtype. The medoid for H5Nx North American sequences was computed from avian strains collected in North America since December 2021, while the control medoid was derived from pre-2020 avian strains. Red lines demarcate the "Mammalian Adaptation Zone," while the distance between the upper red line and the gray line represents the hypothesized "Hard Distance." The first set of boxen plots (left side) shows human-to-avian host-adaptive distances, and the second set (right side) shows non-human mammal-to-avian host-adaptive distances.

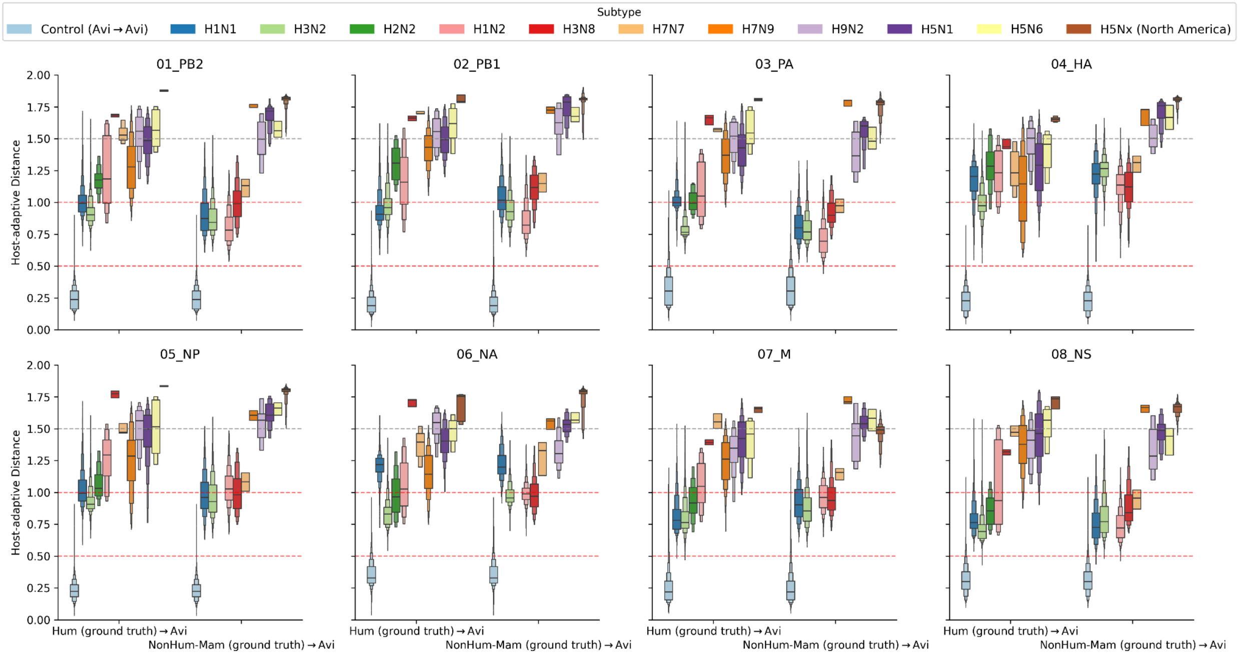

**Supplementary Figure S13. The host-source prediction spectrum of various subtypes.** The stacked bars represent the predicted proportion of each host group as identified by the calibrated model.

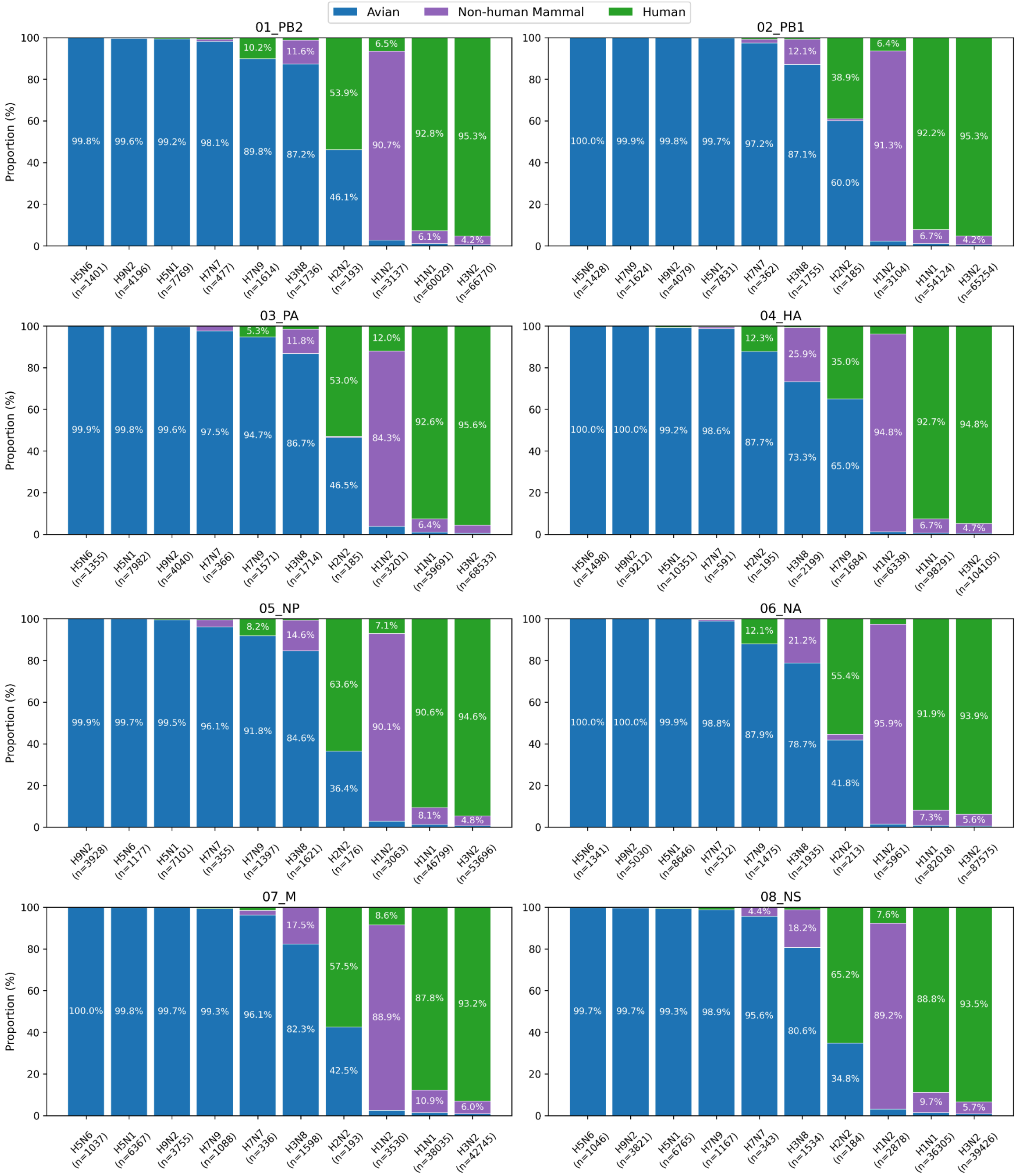

**Supplementary Figure S14. Host-adaptive distances relative to the subtype-specific medoids using avian codon-SHAP profiles.** The boxen plots show human-to-avian, non-human mammal-to-avian, and avian-to-avian (control) host-adaptive distances. These medoids were derived from the codon-SHAP profiles of pre-2020 avian strains within each subtype. The medoid for H5Nx North American sequences was computed from avian strains collected in North America since December 2021, while the control medoid was derived from pre-2020 avian strains. Red lines demarcate the "Mammalian Adaptation Zone." The first set of boxen plots (left side) shows human-to-avian host-adaptive distances, and the second set (right side) shows non-human mammal-to-avian host-adaptive distances.

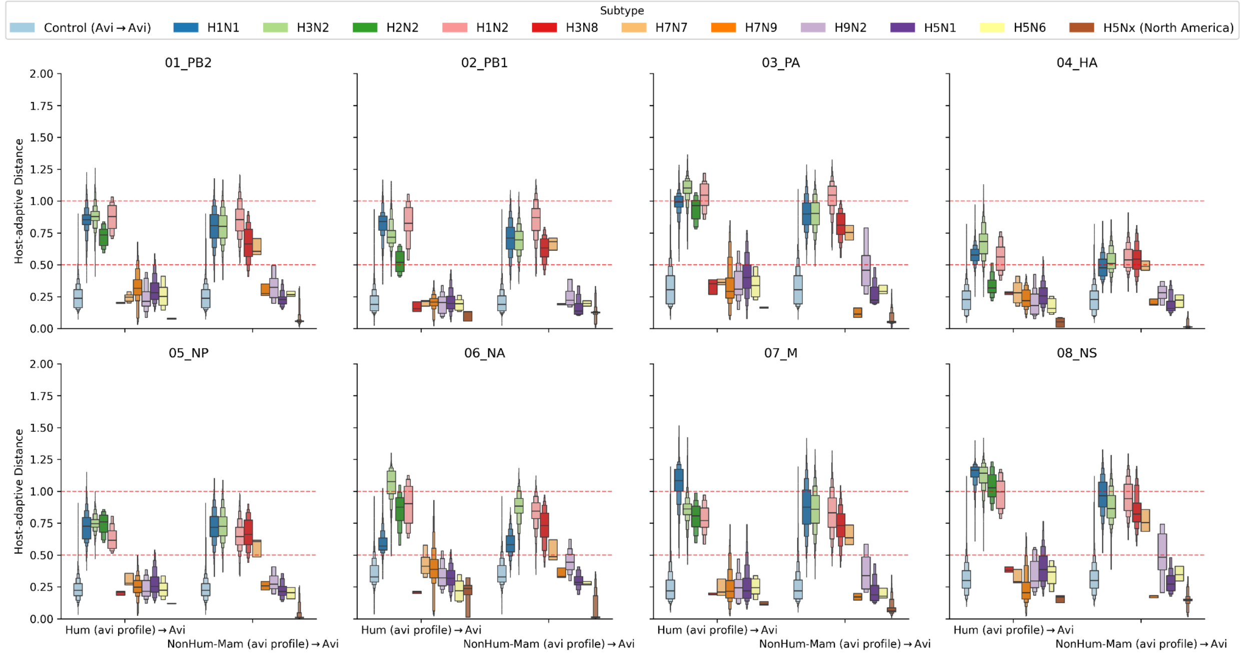

**Supplementary Figure S15: SHAP analysis of non-human mammalian isolate *A/Harbor_Seal/QC/FAV-0836-4/2022* (EPI3053167) showing WaveSeekerNet identifies PB2-E627K.** Panel A shows the global feature importance (SHAP values) for 61 sense codons, ranked from most negative to most positive contributions to the non-human mammalian class. Panel B shows the SHAP values for each nucleotide of the AAG codon at position 627 as a sequence logo of model importance scores. Panel C shows the segment-wide SHAP values of lysine (K), encoded by AAA or AAG, with a clear peak at position 627. Panel D shows the top 20 positively contributing nucleotide positions.

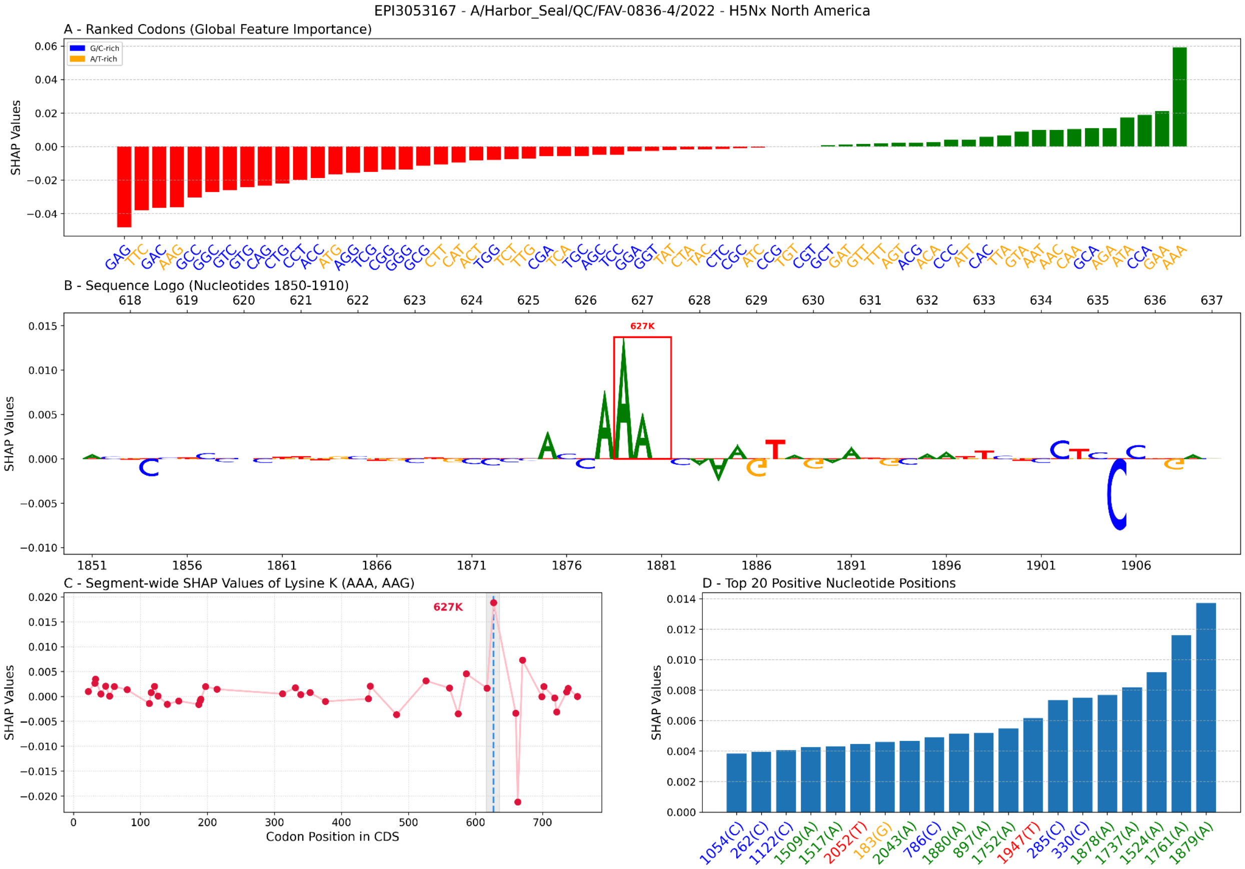

**Supplementary Figure S16: SHAP analysis of the non-human mammalian isolate *A/dairy cow/USA/002644-003/2025* (EPI4097884) showing WaveSeekerNet identifies PB2-D701N.** Panel A shows the global feature importance (SHAP values) for 61 sense codons, ranked from most negative to most positive contributions to the non-human mammalian class. Panel B shows the SHAP values for each nucleotide of the AAC codon at position 701 as a sequence logo of model importance scores. Panel C shows the segment-wide SHAP values of asparagine (N), encoded by AAC or AAT, with a clear peak at position 701. Panel D shows the top 20 positively contributing nucleotide positions.

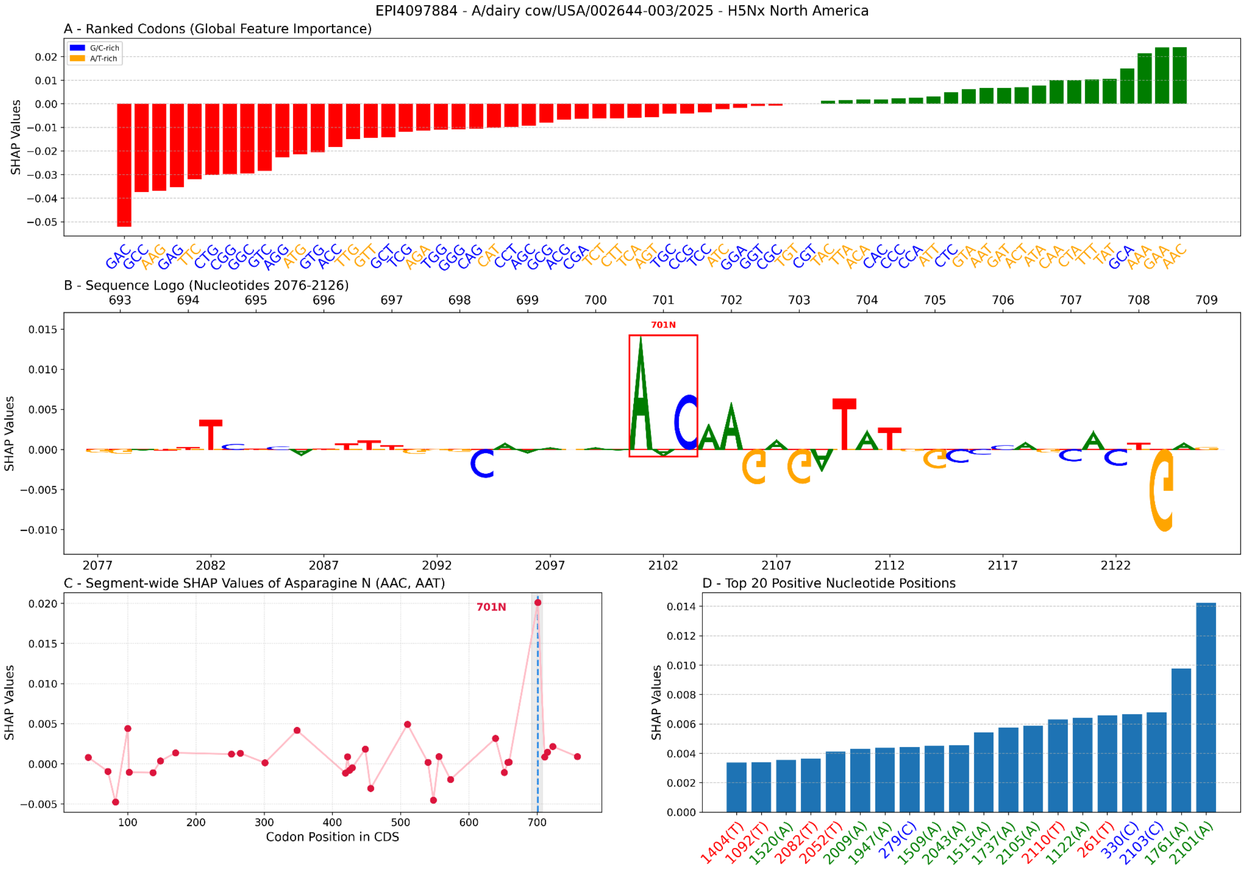
